# Gle1 is required for tRNA to stimulate Dbp5 ATPase activity *in vitro* and to promote Dbp5 mediated tRNA export *in vivo*

**DOI:** 10.1101/2023.06.29.547072

**Authors:** Arvind Arul Nambi Rajan, Ryuta Asada, Ben Montpetit

## Abstract

Cells must maintain a pool of processed and charged transfer RNAs (tRNA) to sustain translation capacity and efficiency. Numerous parallel pathways support the processing and directional movement of tRNA in and out of the nucleus to meet this cellular demand. Recently, several proteins known to control messenger RNA (mRNA) transport were implicated in tRNA export. The DEAD-box Protein 5, Dbp5, is one such example. In this study, genetic and molecular evidence demonstrates that Dbp5 functions parallel to the canonical tRNA export factor Los1. *In vivo* co-immunoprecipitation data further shows Dbp5 is recruited to tRNA independent of Los1, Msn5 (another tRNA export factor), or Mex67 (mRNA export adaptor), which contrasts with Dbp5 recruitment to mRNA that is abolished upon loss of Mex67 function. However, as with mRNA export, overexpression of Dbp5 dominant-negative mutants indicates a functional ATPase cycle and that binding of Dbp5 to Gle1 is required by Dbp5 to direct tRNA export. Biochemical characterization of the Dbp5 catalytic cycle demonstrates the direct interaction of Dbp5 with tRNA (or double stranded RNA) does not activate Dbp5 ATPase activity, rather tRNA acts synergistically with Gle1 to fully activate Dbp5. These data suggest a model where Dbp5 directly binds tRNA to mediate export, which is spatially regulated via Dbp5 ATPase activation at nuclear pore complexes by Gle1.

## INTRODUCTION

Key to the production of proteins is the delivery of amino acids to ribosomes by transfer RNAs (tRNAs). Precursor tRNAs (pre-tRNAs) are first transcribed by RNA Polymerase III with a 5’ leader and a 3’ trailer sequence (Guerrier-Takada, Gardiner et al. 1983, Cook, Fukuhara et al. 2009, Skowronek, Grzechnik et al. 2014, Harris 2016, Graczyk, Ciesla et al. 2018). To function in translation, pre-tRNAs must be end matured and modified, spliced, amino-acylated and exported from the nucleus in eukaryotes. Importantly, it has come to be appreciated that these steps in tRNA processing are not necessarily sequential (Trotta, Miao et al. 1997, Yoshihisa, Yunoki-Esaki et al. 2003, Yoshihisa, Ohshima et al. 2007, Wu and Hopper 2014, Hopper and Huang 2015, Wu, Bao et al. 2015, Phizicky and Hopper 2023). For example, in the budding yeast *Saccharomyces cerevisiae,* intron-containing tRNAs are spliced in the cytoplasm on the mitochondrial surface by the tRNA Splicing Endonuclease (SEN) complex following nuclear export (Yoshihisa, Yunoki-Esaki et al. 2003, Yoshihisa, Ohshima et al. 2007, Skowronek, Grzechnik et al. 2014, Wan and Hopper 2018). Furthermore, cytoplasmic tRNAs can be reimported to the nucleus through retrograde transport to allow tRNAs to undergo further modification and quality control which is also used to repress translation during conditions of stress (Murthi, Shaheen et al. 2010, Hopper and Huang 2015, Nostramo and Hopper 2020). As tRNAs cycle between the nucleus and cytoplasm, they undergo various chemical modifications that are also spatially regulated (Phizicky and Hopper 2023). The proper processing and modification of tRNAs through this complex regulatory network dictates proper folding of pre-tRNAs and recognition by appropriate tRNA binding proteins that control subsequent steps in the tRNA life cycle.

In the context of nuclear export, end maturation and amino-acylation are required for recognition by the tRNA Exportin Los1/Exportin-t and Msn5 respectively (Huang and Hopper 2015, Chatterjee, Majumder et al. 2017, Nostramo and Hopper 2020, Chatterjee, Marshall et al. 2022). Although Los1 and Msn5 are the best characterized tRNA export factors, both are non-essential and can be deleted in combination (Takano, Endo et al. 2005, Murthi, Shaheen et al. 2010). Given the essential nature of tRNA export, this has prompted several studies, including a comprehensive screen of nearly all annotated genes in *S. cerevisiae,* to identify additional factors responsible for regulating nucleocytoplasmic dynamics of tRNA localization and processing (Feng and Hopper 2002, Steiner-Mosonyi, Leslie et al. 2003, Eswara, McGuire et al. 2009, Huang and Hopper 2015, Wu, Bao et al. 2015, Chatterjee, Majumder et al. 2017, Nostramo and Hopper 2020, Chatterjee, Marshall et al. 2022). These studies have elucidated functional roles of the karyopherin Crm1 and mobile nucleoporin Mex67 in regulating tRNA nucleocytoplasmic dynamics (Chatterjee, Majumder et al. 2017, Derrer, Mancini et al. 2019, Nostramo and Hopper 2020, Chatterjee, Marshall et al. 2022). Both factors have previously been shown to function in messenger RNA (mRNA) export (Hodge, Colot et al. 1999, Takemura, Inoue et al. 2004, Lund and Guthrie 2005, Derrer, Mancini et al. 2019), with Mex67 serving an essential role in the process (Hodge, Colot et al. 1999, Lund and Guthrie 2005, Adams and Wente 2020). In tRNA export, it is thought that Crm1 and Mex67 both function parallel to Los1 to support export of pre-tRNA and re-export of spliced tRNA that have undergone retrograde transport (Chatterjee, Majumder et al. 2017, Nostramo and Hopper 2020, Chatterjee, Marshall et al. 2022). Moreover, recent reports suggest subspecies specialization and family preferences for tRNA cargo, with Mex67 serving a unique function in the premature export of pre-tRNAs that have not completed 5’ end processing (Chatterjee, Marshall et al. 2022).

While non-coding RNA (ncRNA) and mRNA export pathways have long been hypothesized to be regulated by independent processes, recent studies indicate the participation of numerous mRNA export factors in regulation of diverse ncRNAs (Yao, Roser et al. 2007, Yao, Lutzmann et al. 2008, Faza, Chang et al. 2012, Bai, Moore et al. 2013, Wu, Becker et al. 2014, Becker, Hirsch et al. 2019, Vasianovich, Bajon et al. 2020). In addition to tRNA and mRNA, both Mex67 and Crm1 have functions in pre-snRNA, pre-ribosomal subunit, and *TLC1* (telomerase) export (Yao, Roser et al. 2007, Faza, Chang et al. 2012, Bai, Moore et al. 2013, Wu, Becker et al. 2014, Becker, Hirsch et al. 2019, Vasianovich, Bajon et al. 2020). Similarly, in addition to Mex67 and Crm1, the DEAD-box Protein 5 (Dbp5) has also been implicated in each of these RNA export pathways, including tRNA export (Wu, Becker et al. 2014, Neumann, Wu et al. 2016, Mikhailova, Shuvalova et al. 2017, Becker, Hirsch et al. 2019, Lari, Arul Nambi Rajan et al. 2019). Dbp5 is most well known as an RNA-stimulated ATPase that is spatially regulated by the nucleoporins Nup159 and Gle1 with the small molecule inositol hexakisphosphate (InsP_6_) at the cytoplasmic face of the nuclear pore complex (NPC) to promote the directional export of mRNA (Snay-Hodge, Colot et al. 1998, Tseng, Weaver et al. 1998, Hodge, Colot et al. 1999, Schmitt, von Kobbe et al. 1999, Strahm, Fahrenkrog et al. 1999, Zhao, Jin et al. 2002, Weirich, Erzberger et al. 2004, Lund and Guthrie 2005, Alcazar-Roman, Tran et al. 2006, Weirich, Erzberger et al. 2006, Tran, Zhou et al. 2007, Dossani, Weirich et al. 2009, Fan, Cheng et al. 2009, von Moeller, Basquin et al. 2009, Hodge, Tran et al. 2011, Montpetit, Thomsen et al. 2011, Noble, Tran et al. 2011, Kaminski, Siebrasse et al. 2013, Wong, Cao et al. 2016, Adams, Mason et al. 2017, Wong, Gray et al. 2018, Adams and Wente 2020). In addition to these export roles, Dbp5 also shuttles between the nucleus and cytoplasm with reported roles in transcription, R-loop metabolism, and translation (Hodge, Colot et al. 1999, Estruch and Cole 2003, Gross, Siepmann et al. 2007, Scarcelli, Viggiano et al. 2008, Tieg and Krebber 2013, Mikhailova, Shuvalova et al. 2017, Lari, Arul Nambi Rajan et al. 2019, Beissel, Grosse et al. 2020). Notably, Nup159 and Gle1 mutants also alter tRNA export (Lari, Arul Nambi Rajan et al. 2019), suggesting the mRNA export pathway as a whole (e.g., Mex67, Dbp5, Gle1/ InsP_6_, Nup159) may support tRNA export. However, a mechanistic understanding of how Dbp5 supports these diverse functions and whether they are regulated by similar co-factors and enzymatic activity is largely not understood.

A recent mutagenesis screen within our research group identified mutants that alter the nucleocytoplasmic shuttling dynamics of Dbp5 (Lari, Arul Nambi Rajan et al. 2019). Mutations were identified (e.g., *dbp5^L12A^*) that disrupt a nuclear export sequence (NES) recognized by Crm1 to promote Dbp5 transport out of the nucleus. In contrast, another mutant, *dbp5^R423A^*, was found to be impaired in nuclear import from the cytoplasm. Importantly, neither of these mutations disrupt the essential mRNA export functions of Dbp5; however, limited nuclear access of *dbp5^R423A^* induced tRNA export defects suggesting a potentially novel role for nuclear Dbp5 in tRNA export. In support of this hypothesis, a physical interaction between Dbp5 and tRNA was also characterized, which was elevated upon nuclear accumulation of Dbp5 in a *dbp5^L12A^*mutant (Lari, Arul Nambi Rajan et al. 2019).

In this study, genetic and biochemical characterization of Dbp5 mediated tRNA export was performed. The data show that Dbp5 is recruited to pre-tRNA independent of Los1 and Msn5, indicating Dbp5 has functions independent of the primary Los1 mediated pre-tRNA export pathway. Similarly, Dbp5 does not require Mex67 as an adapter for tRNA binding (unlike proposed mechanisms of mRNA export). In contrast to single stranded RNA substrates (e.g., mRNA), this study further demonstrates an interaction of Dbp5 with tRNA and double stranded RNA (dsRNA) that leads to robust ATPase activation only in the presence of Gle1/ InsP_6_. These findings, together with previous research (Lari, Arul Nambi Rajan et al. 2019), suggest that Dbp5 engages tRNA within the nucleus in a manner distinct from mRNA, and requires Gle1/ InsP_6_ to mediate RNA stimulated ATPase activation of Dbp5 to promote pre-tRNA export.

## RESULTS

### Dbp5 has functions independent of Los1 in pre-tRNA export

Previous studies have placed the mRNA export factor Mex67, a proposed target of Dbp5 regulation in mRNA export (Lund and Guthrie 2005, Adams and Wente 2020), in a pathway parallel to Los1 regulated pre-tRNA nucleo-cytoplasmic shuttling (Chatterjee, Majumder et al. 2017). To investigate the relationship between Dbp5 and Los1, it was first determined if the subcellular distribution of either protein was dependent on the activity of the other. In a *los1τι,* the NPC localization of GFP-Dbp5 was not altered (Figure 1A). Similarly, when a Dbp5 loss of function was induced using Auxin Induced Degradation (AID) (Nishimura, Fukagawa et al. 2009), Los1-GFP localization remained unchanged (Figure 1B) while mRNA export defects were detected (Supplement 1A), confirming depletion of Dbp5 to a level sufficient to disrupt mRNA export. To test the genetic relationship of Los1 and Dbp5, a *los1τι* was combined with the previously identified Dbp5 shuttling mutants (i.e., *dbp5^L12A^* and *dbp5^R423^*). Double mutants of *los11ι* with *dbp5^L12A^* (increased in the nucleus) or *dbp5^R423^*(depleted from the nucleus) exhibited no growth defects (Figure 1C). Similarly, combining *dbp5^L12A^* and *dbp5^R423A^* respectively with *los1Δ/msn5Δ* double mutants (Supplement 1B) revealed minimal growth defects. The viability of these mutants suggests Dbp5 may support Los1 and Msn5 tRNA export pathways or the existence of a parallel pathway(s) that is able to maintain cellular fitness in the presence of *dbp5^R423A^*. Importantly, this genetic relationship between *dbp5^R423A^* and *los11ι* mirrors those previously published for *mex67-5* and *los11ι* strain (Chatterjee, Majumder et al. 2017). To further distinguish these possibilities, tRNA FISH was performed using a probe specific to the pre-tRNA^Ile^_UAU_ intron (SRIM03) in single and double mutants. The *los11ι dbp5^R423A^* double mutant exhibited significantly stronger nuclear accumulation of pre-tRNA as compared to single mutants or control after a 4-hour incubation at 37°C (Figure 1D/E). Notably, elevating nuclear pools of Dbp5 by combining *dbp5^L12A^* with *los11ι* did not alter the pre-tRNA export defects observed in *los11ι* strains, indicating that excess nuclear Dbp5 is not sufficient to suppress the *los11ι* phenotype. These results were recapitulated with FISH probes targeting precursor/mature isoforms of tRNA^Ile^_UAU_ and tRNA^Tyr^_GUA_ (Supplement 1C/D). These data argue for a role for Dbp5 in pre-tRNA export that operates independently of Los1.

**Figure 1:**
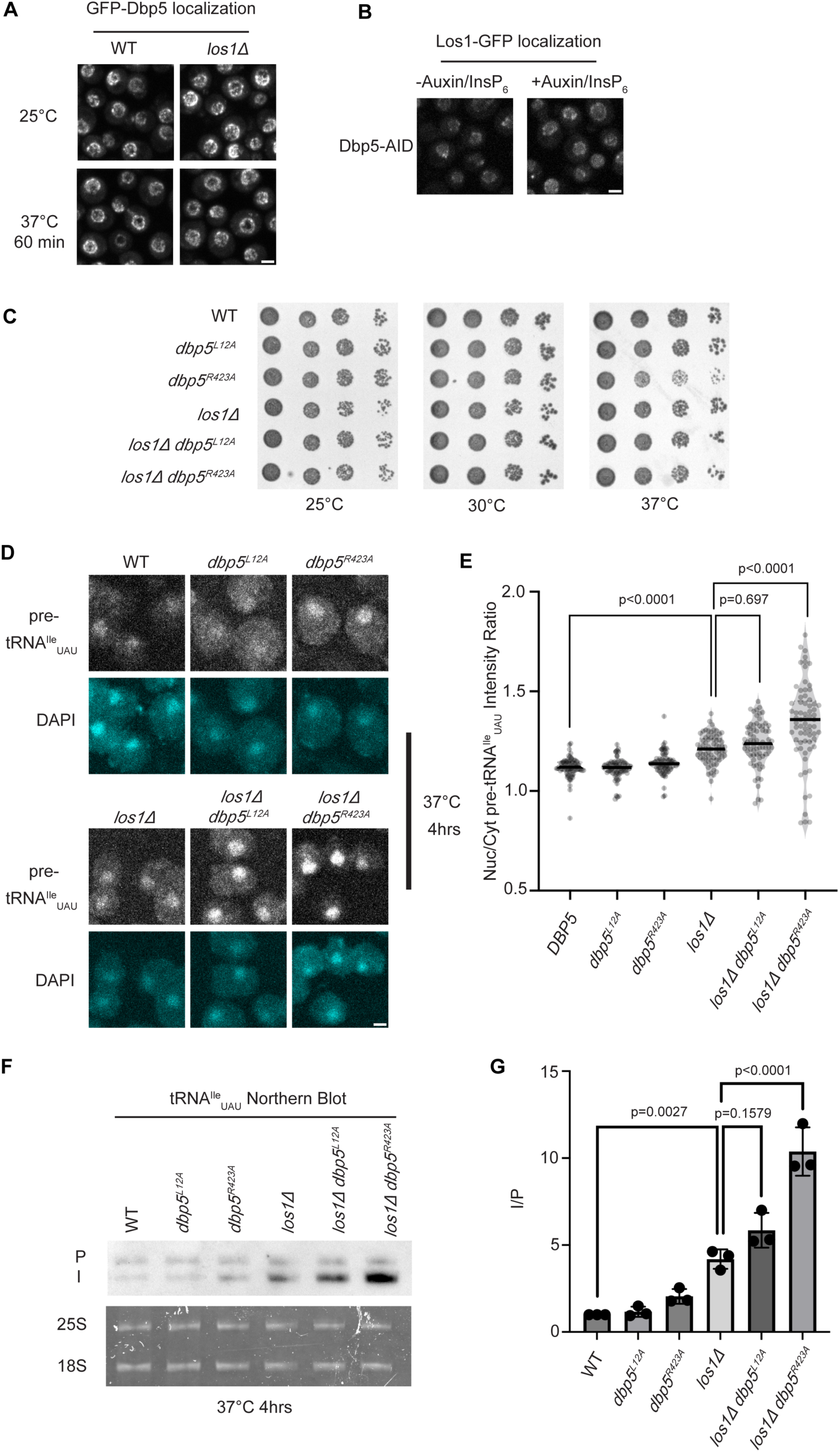
Dbp5 functions parallel to Los1 in pre-tRNA export. (A) Fluorescent images show GFP-Dbp5 remains enriched at the nuclear periphery in wild-type and *los111* at 25°C and 37°C. Scale bar represents 2 µm. (B) Los1-GFP remains nucleoplasmic and associated with nuclear periphery in Dbp5-AID after treatment with DMSO or 500 µM Auxin and 10 µM InsP_6_ for 90 minutes. Scale bar represents 2 µm. (C) Spot assay for growth of strains containing untagged *dbp5^L12A^, dbp5^R423A^,* or *los111* integrated at the endogenous gene locus after two days at 25, 30, and 37°C on YPD. (D) tRNA FISH targeting intron of tRNA^Ile^_UAU_ in indicated strains after pre-culture to early log phase at 25°C and shift to 37°C for 4 hours. Scale bar represents 2 µm. (E) Quantification of tRNA FISH from (E). Ratio of average nuclear to cytoplasmic pixel intensities were calculated across 3 independent replicate experiments and pooled for plotting. P-values were calculated using one-way ANOVA. (F) Northern Blot analysis targeting precursor and mature isoforms of tRNA^Ile^_UAU_. Small RNAs were isolated from strains at mid log phase growth after pre-culture at 25°C and shift to 37°C for 4 hours. “P” bands represent intron containing precursors that have 5’ leader/3’ trailer sequences and “I” bands represent intron-containing end processed tRNA intermediates that have leader/trailer sequences removed. (G) Quantification of Northern blot from (G). Ratio of signal from intron-containing end processed intermediates (I) vs 5’ leader/3’ trailer containing precursor (P) was calculated and presented relative to I/P ratio observed for WT. Error bars represent standard deviation and p-values calculated using one-way ANOVA.

To complement the FISH data, Northern blotting analyses were conducted to assess tRNA processing. Intron-containing pre-tRNAs are transcribed with 5’ leader and 3’ trailer sequences to generate precursor molecules (P) that are end-processed in the nucleus to generate intron-containing intermediates (I) that can be detected by Northern blotting (Wu, Bao et al. 2015). When nuclear export of the end-processed pre-tRNA is disrupted a change in the precursor (P) to intron-containing intermediate (I) ratio is observed (Wu, Huang et al. 2013, Wu, Bao et al. 2015, Chatterjee, Majumder et al. 2017). As such, I/P ratios were measured in the single and *los11ι/dbp5* double mutants. In line with the increased nuclear localization of pre-tRNAs observed by FISH, an additive defect in pre- tRNA^Ile^_UAU_ and pre- tRNA^Tyr^_GUA_ was observed when *los11ι* was combined *dbp5^R423A^* (∼2-fold increase in the I/P ratio relative to *los11ι* alone, Figure 1F/G and Supplement 1E/F). These FISH and Northern blotting data mirror phenotypes previously reported for mutants of Mex67 in relation to Los1 (Chatterjee, Marshall et al. 2022), and indicate that Dbp5 has functions in pre-tRNA export that are independent of Los1.

### Known tRNA export factors do not recruit Dbp5 to pre-tRNAs in vivo

Dbp5, like other DEAD-box proteins, has been reported to bind nucleic acids through sequence independent interactions with the phosphate backbone (Andersen, Ballut et al. 2006, Bono, Ebert et al. 2006, Sengoku, Nureki et al. 2006, Montpetit, Thomsen et al. 2011). For other helicases of the SF2 family, specificity in RNA substrates is conferred by adaptor proteins (Lund and Guthrie 2005, Jankowsky 2011, Thoms, Thomson et al. 2015). For this reason, it was investigated whether known tRNA export factors aid recruitment of Dbp5 to pre-tRNA substrates. To do so, RNA immunoprecipitation (RIP) experiments were performed with protein-A (prA) tagged Dbp5, integrated at its endogenous locus and present as the sole copy of the gene, in strains where tRNA export factors were deleted. Co-immunoprecipitated RNAs were analyzed by RT-qPCR with primers specific to un-spliced intron-containing pre-tRNA^Ile^_UAU._ The abundance of the pre-tRNA^Ile^_UAU_ target in each IP was normalized to the abundance of the target in the corresponding input sample to control for changes in gene expression. Relative enrichment of the target RNA was then compared to the background signal obtained from RNA IPs in a common untagged control. In a *los11ι* strain, a ∼2-fold reduction in the relative amount of pre-tRNA co-immunoprecipitated with Dbp5 (∼9.5-fold enrichment in WT to ∼4.5-fold in *los11ι*) was observed, but importantly the deletion failed to abolish the Dbp5 pre-tRNA *in vivo* interaction (Figure 2A). Loss of Msn5, which does not have a published role in the export of intron-containing pre-tRNAs (Huang and Hopper 2015), did not cause a significant change in Dbp5 pre-tRNA interactions by RNA IP. These results indicate that Los1 may support Dbp5-pre-tRNA complex formation, but Dbp5 maintains an ability to bind pre-tRNA *in-vivo* in the absence of Los1. This finding is consistent with the additive tRNA export defects observed in *los11ι dbp5^R423A^* double mutants (Figure 1) and support the hypothesis that Dbp5 has a function in tRNA export parallel to Los1.

**Figure 2:**
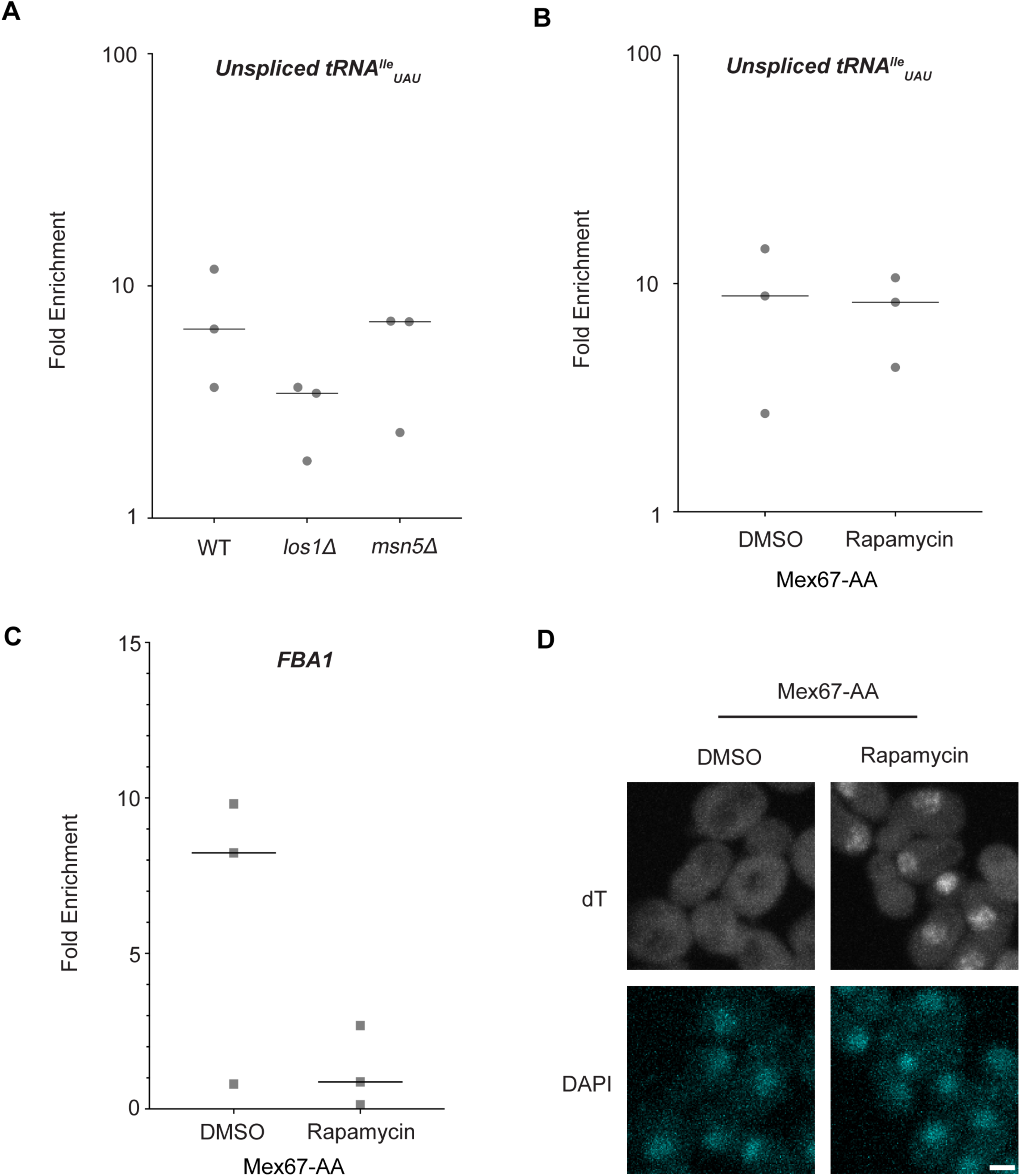
Los1 and Mex67 are not required for Dbp5 recruitment to pre-tRNA^Ile^_UAU_. (A) Plots show relative fold enrichment of tRNA^Ile^_UAU_ following prA-Dbp5 RNA IP in a wild-type (WT), *los111,* and *msn511* strain background. Abundance of target gene is normalized to abundance of the target transcript in input samples and represented as fold enrichment relative to RNA IP from an untagged control. (B) prA-Dbp5 RNA IP targeting tRNA^Ile^_UAU_ in Mex67-AA after either treatment with DMSO or 1 µg/ml rapamycin for 15 minutes. (C) prA-Dbp5 RNA IP targeting FBA1 mRNA in Mex67-AA after either treatment with DMSO or 1 µg/ml rapamycin for 15 minutes. (D) dT FISH confirming a mRNA export defect caused by loss of function of Mex67 following 15min incubation with 1 µg/ml rapamycin compared to a DMSO control. Scale bar represents 2 µm.

In mRNA export, Mex67 is a proposed target of Dbp5 activity to promote directional nuclear export (Adams and Wente 2020). Since neither of the non-essential tRNA export factors abolished Dbp5-tRNA interaction when deleted, it was tested if Mex67 has a role in recruiting Dbp5 to tRNA. An Anchors Away approach was used to rapidly re-localize Mex67 to the cytoplasmic peroxisomal anchor (Pex25-FKBP12) in an inducible manner dependent on the addition of rapamycin (Haruki, Nishikawa et al. 2008). The resulting strains were sensitive to rapamycin, causing nuclear mRNA accumulation by dT FISH within 15 minutes of addition (Fig 2D). This timing is consistent with a previous report employing Mex67 Anchors Away (Mex67-AA) (Tudek, Schmid et al. 2018). PrA-Dbp5 was integrated at its endogenous locus in these strains and RNA-IP experiments were performed before and after rapamycin addition as described above. Under these conditions of Mex67-AA re-localization, no significant change in the Dbp5-tRNA^Ile^_UAU_ interaction was observed (median of ∼8.8-fold enrichment before addition of rapamycin and ∼8.2-fold after, Figure 2B). Importantly, consistent with the function of Mex67 in mRNA export (Snay-Hodge, Colot et al. 1998, Tseng, Weaver et al. 1998, Schmitt, von Kobbe et al. 1999, Strahm, Fahrenkrog et al. 1999, Weirich, Erzberger et al. 2004, Lund and Guthrie 2005, Tran, Zhou et al. 2007, Dossani, Weirich et al. 2009, von Moeller, Basquin et al. 2009, Adams and Wente 2020), the nuclear retention of mRNAs via Mex67-AA resulted in strongly reduced binding of Dbp5 to the *FBA1* mRNA (from a median of ∼8 fold enrichment above background before addition of rapamycin to no enrichment after, Figure 2C). Together, these RNA IP experiments suggest that Dbp5 binds to tRNAs independent of known tRNA export proteins and does so in a manner that is distinct from mRNA.

### The Dbp5 ATPase cycle supports tRNA export in vivo

In mRNA export, RNA binding, Gle1 activation, and ATP hydrolysis by Dbp5 are all critical for directional transport (Hodge, Tran et al. 2011, Noble, Tran et al. 2011). The *dbp5^R426Q^* and *dbp5^R369G^*mutants have previously been shown to exhibit deficient RNA binding (<5% of WT), ATPase activity (∼10% and 60% of WT, respectively) and have a dominant-negative effects on mRNA export status when overexpressed (Hodge, Tran et al. 2011, Noble, Tran et al. 2011). Additionally, it has been shown that *dbp5^R369G^* has elevated Gle1 affinity and effectively competes with WT Dbp5 for Gle1 binding (Hodge, Tran et al. 2011). In contrast, *dbp5^E240Q^* is an ATP hydrolysis mutant that is competent to bind RNA, as it is biased to the ATP bound state, but is recessive lethal showing no mRNA export defect when overexpressed in the context of endogenously expressed WT Dbp5 (Hodge, Tran et al. 2011). The *dbp5^R426Q^*and *dbp5^R369G^* mutants localize to the nuclear periphery similar to wild-type Dbp5, however *dbp5^E240Q^*is predominantly cytoplasmic (with some nuclear envelope localization detectable when wild-type Dbp5 is depleted) (Hodge, Tran et al. 2011). Unlike the impact of these mutants on mRNA export, it has been reported that the ability of Dbp5 to complete ATP hydrolysis (but not RNA binding) is dispensable for pre-ribosomal subunit export, as is Gle1 (Neumann, Wu et al. 2016).

Based on the observation that Gle1 functions to support tRNA export (Lari, Arul Nambi Rajan et al. 2019), these Dbp5 mutants deficient in ATPase activity or altered Gle1 binding were tested for their influence on tRNA export to infer if tRNA export is more similar to mRNA or pre-ribosomal subunit export. To individually express these lethal mutants and the wildtype control, *DBP5*, *dbp5^R426Q^, dbp5^R369G^,* or *dbp5^E240Q^*were integrated at the *URA3* locus under regulation of the inducible pGAL promoter. Protein expression was induced for 6 hours by shifting cells from raffinose to galactose containing media. Induction was followed by 1 hour of growth in glucose to halt expression and relieve potential changes in tRNA export induced by a change in the primary carbon source. Using this approach, all proteins were expressed as indicated by western blotting (Figure 3A) and showed the previously reported impact on mRNA export (Figure 3B). Northern blotting targeting tRNA^Ile^_UAU_ revealed both *dbp5^R426Q^*and *dbp5^R369G^,* but not *dbp5^E240Q^,* showed an accumulation of intron-containing precursor tRNA^Ile^_UAU_ as compared to over expression of wildtype Dbp5 (I/P ratio of 1.97 +/- 0.06 vs 1.93 +/- 0.31 vs 1.32 +/- 0.41, respectively) (Figure 3C/D). These phenotypes differ from the reported impacts on pre-ribosomal subunit transport, indicating Dbp5 functions differently in these non-coding RNA export pathways. Furthermore, these data indicate that both the ATPase cycle and regulation of Dbp5 by Gle1 are central to the function of Dbp5 in tRNA export, as each activity is in mRNA export.

**Figure 3:**
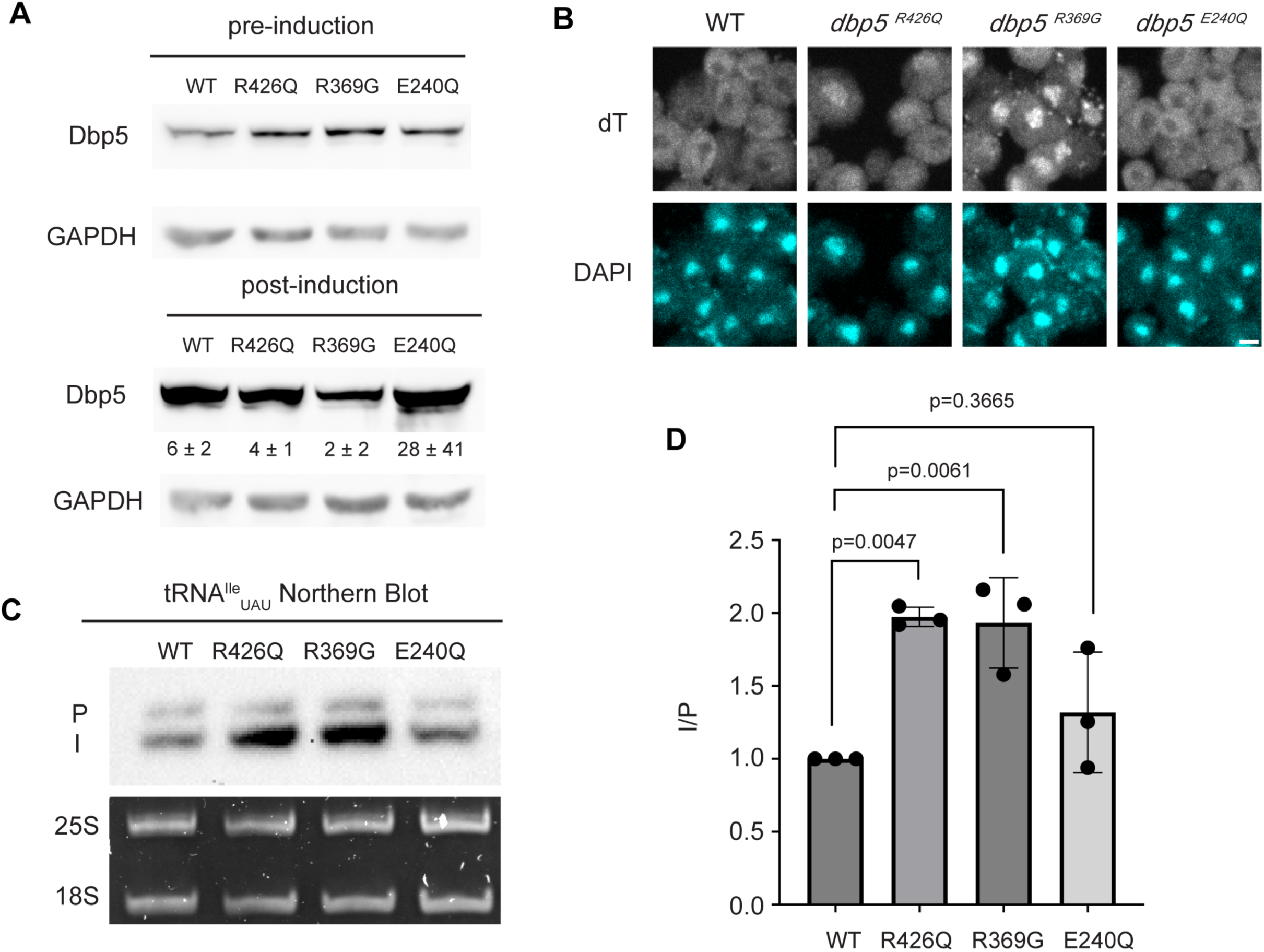
ATPase activity and Gle1 is required for Dbp5-mediated tRNA export in vivo. (A) Western blot with either mouse monoclonal anti-DBP5 or anti-GAPDH (loading control) antibody to detect overexpression of untagged Dbp5 and Dbp5 ATPase mutants. Strains were cultured in raffinose at 25°C to derepress pGAL promoter (pre-induction) and expression was induced by shifting cultures into 2% galactose containing media for 6 hours (post-induction). Expression was stopped by addition of glucose for 1 hour. Overexpression was observed over the level of wildtype Dbp5 that is expressed from the endogenous locus and is present in the pre-induction samples. Top and bottom panels are from the same blot and processed equally. Quantification of the extent overexpression in post-induction samples is presented below Dbp5 bands. Signal intensity of Dbp5 bands were normalized to abundance of Gapdh and presented relative to pre-induction levels of expression. (B) dT FISH confirms previously reported mRNA export status phenotypes for Dbp5 ATPase mutants with *dbp5^R426Q^* and *dbp5^R369G^* showing a nuclear accumulation of poly(A)-RNA. Scale bar represents 2 µm. (C) Northern blot analysis targeting precursor and mature isoforms of tRNA^Ile^_UAU_ from yeast strains overexpressing *DBP5*, *dbp5^R426Q^, dbp5^R369G^,* or *dbp5^E240Q^*. (D) Quantification of Northern blot from (C). Ratio of signal from intron-containing end processed intermediates (I) vs 5’ leader/3’ trailer containing precursor (P) was calculated and presented relative to I/P ratio observed for WT. Error bars represent standard deviation and p-values calculated using one-way ANOVA.

### Dbp5 ATPase activity in the presence of tRNA is Gle1-dependent

Given the impact of *dbp5^R426Q^* and *dbp5^R369G^*overexpression on tRNA export *in vivo*, the Dbp5 ATPase cycle and role of Gle1 in the context of tRNA was further investigated *in vitro*. Previous studies have characterized Dbp5 binding to single stranded RNA (ssRNA) substrates and demonstrated Dbp5 binding to ssRNA is linked to the nucleotide state of the enzyme, with the highest affinity for ssRNA occurring when Dbp5 is ATP bound (Weirich, Erzberger et al. 2006, Montpetit, Thomsen et al. 2011, Arul Nambi Rajan and Montpetit 2021). Following ssRNA binding, hydrolysis of ATP to ADP results in a reduction in the affinity of Dbp5 for the ssRNA substrate (Weirich, Erzberger et al. 2006). While there has been extensive characterization of the structural and biochemical details of Dbp5, ssRNA, and co-regulators (Wong, Gray et al. 2018, Mason and Wente 2020, Gray, Cao et al. 2022), no such understanding exists for Dbp5 engaging highly structured RNA substrates like tRNA. As such, it was first tested if recombinant full length Dbp5 could bind commercially available yeast mixed tRNAs or the yeast Phenylalanine tRNA (both substrates that have been used in previous biochemical and structural studies (Shi and Moore 2000, Yao, Roser et al. 2007)).

To test whether Dbp5 can form complexes with tRNA *in vitro,* electrophoretic mobility shift assays (EMSA) were performed in the presence of different nucleotide states and tRNA substrates. Consistent with published observations of complex formation between Dbp5 and ssRNA (Weirich, Erzberger et al. 2006, Montpetit, Thomsen et al. 2011), Dbp5 generated band shifts for both mixed tRNA substrates and Phe tRNA in the presence of the ATP state analog ADP•BeF_3_ (Figure 4A). In the presence of ATP or ADP, or with no nucleotide, Dbp5 failed to form a similar band shift, which exactly parallels reported interactions between Dbp5 and ssRNA in these nucleotide states (Weirich, Erzberger et al. 2006, Montpetit, Thomsen et al. 2011).

**Figure 4:**
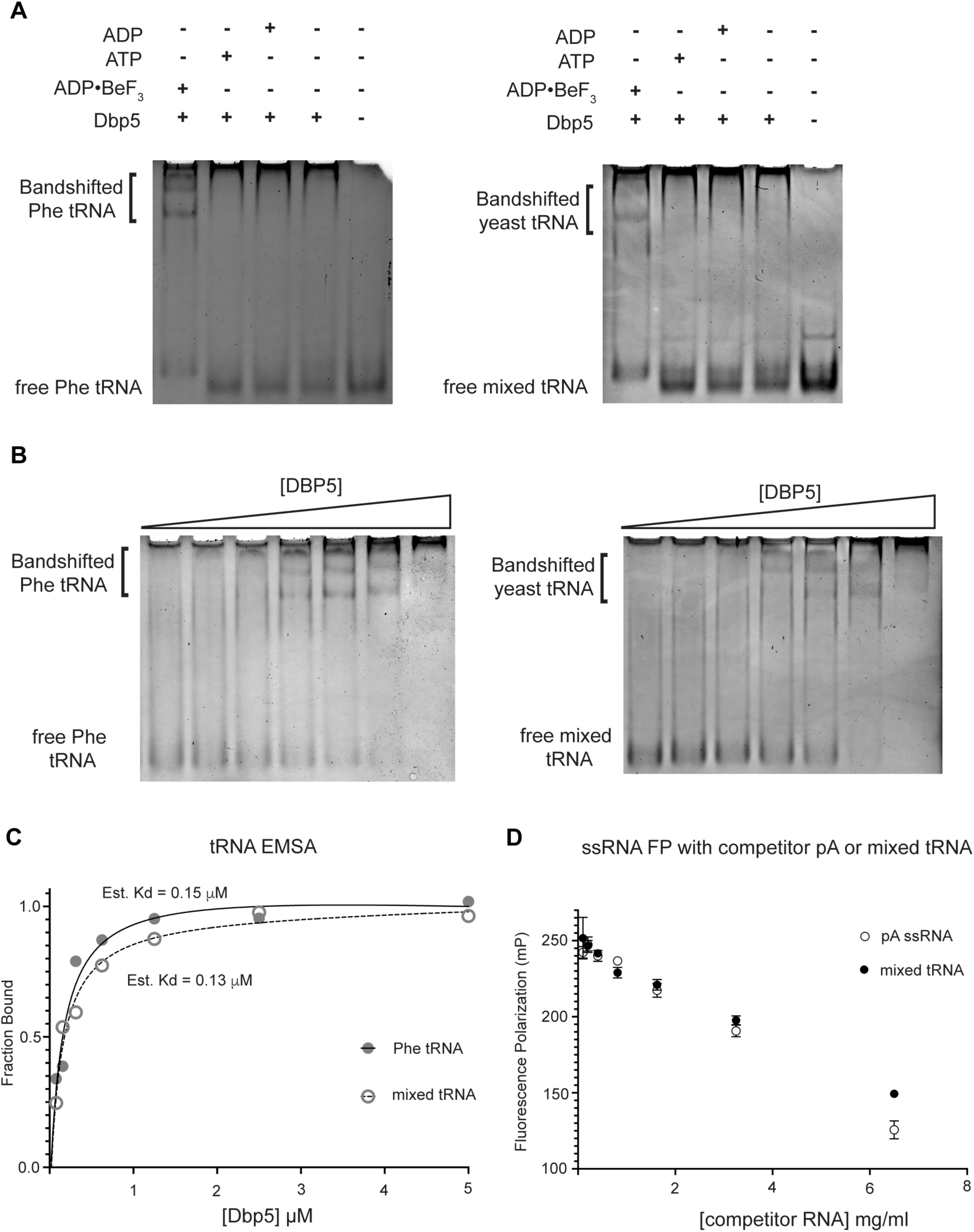
Dbp5 binds yeast tRNA in vitro. (A) Full length recombinant Dbp5 (fl-Dbp5) binds both mixed yeast tRNA and Phenylalanine (Phe) tRNA in the presence of the ATP mimetic ADP•BeF_3_ by electrophoresis mobility shift assay (EMSA). RNA binding reaction was conducted in the presence of ADP•BeF_3_, ATP, ADP, or no nucleotide and resolved on 6% native polyacrylamide gel at 70V. Reactions contained 2 µM fl-Dbp5, 1 mM nucleotide when present, 250ng tRNA and binding buffer. (B) EMSA experiments in which increasing concentrations of fl-Dbp5 (1:2 dilution series starting at 5uM Dbp5) was titrated into RNA binding reactions in the presence of 1 mM ADP•BeF_3_, 250ng mixed yeast tRNA or Phe tRNA, and binding buffer. (C) Band intensities of free probe and band shifts were quantified, and bound fraction was calculated for each well of EMSAs in (B). One site binding model was fit to the data using GraphPad Prism to estimate Kd for both mixed yeast tRNA and Phe tRNA. (D) Unlabeled pA ssRNA and mixed yeast tRNA compete with a 16nt fluorescein labelled ssRNA for fl-Dbp5 binding. Fluorescence polarization competition assays were performed by titrating increasing concentration of an unlabeled competitor, pA (open square) or mixed yeast tRNA (closed circle), in reactions containing 50 nM fluorescein labelled ssRNA, 2.5 mM ADP•BeF_3_, 1 µM Dbp5 and buffer. Error bars represent standard deviation of three independent experiments.

To further estimate an affinity for tRNA, both the Phenylalanine tRNA and yeast mixed tRNAs were used in EMSA experiments with varying concentrations of Dbp5 (and fixed concentrations of ADP•BeF_3_ and tRNA). From quantification of EMSAs, an estimated K_d_ of ∼150nM (Phe tRNA) and ∼130nM (mixed tRNA) was obtained (Figure 4B/C). In addition to EMSAs, Dbp5 binding to tRNA was observed and tested in competition assays using fluorescence polarization measurements. In these assays, increasing concentrations of unlabeled tRNA or a ssRNA homopolymeric polyadenylic acid (pA, routinely used for RNA activation in steady-state ATPase assays (Weirich, Erzberger et al. 2006)), were found to compete with a fluorescently labeled ssRNA for Dbp5 in the presence of ADP•BeF_3_ (Figure 4D). These data, from orthogonal approaches, indicate that Dbp5 interacts directly with tRNA *in vitro,* and does so in a manner governed by principles that resemble Dbp5-ssRNA binding.

Interaction with ssRNA substrates has been well documented to stimulate the Dbp5 ATPase cycle (Weirich, Erzberger et al. 2004, Lund and Guthrie 2005, Alcazar-Roman, Tran et al. 2006, Weirich, Erzberger et al. 2006, Tran, Zhou et al. 2007, Dossani, Weirich et al. 2009, von Moeller, Basquin et al. 2009, Hodge, Tran et al. 2011, Montpetit, Thomsen et al. 2011, Noble, Tran et al. 2011). As such, it was next determined if binding of tRNA to Dbp5 stimulates ATPase activity using a spectrophotometric ATPase assay (Weirich, Erzberger et al. 2006, Montpetit, Thomsen et al. 2011). Unlike a ssRNA substrate (i.e., pA) that can maximally stimulate ATP turnover to ∼0.5 ATP/sec, neither tRNA nor poly(I:C) (used as an orthogonal dsRNA substrate) stimulated Dbp5 ATPase activity over a range of RNA concentrations (Figure 5A). These results with the above binding data suggest Dbp5 can engage structured and dsRNA substrates, but this does not lead to productive ATP hydrolysis.

**Figure 5:**
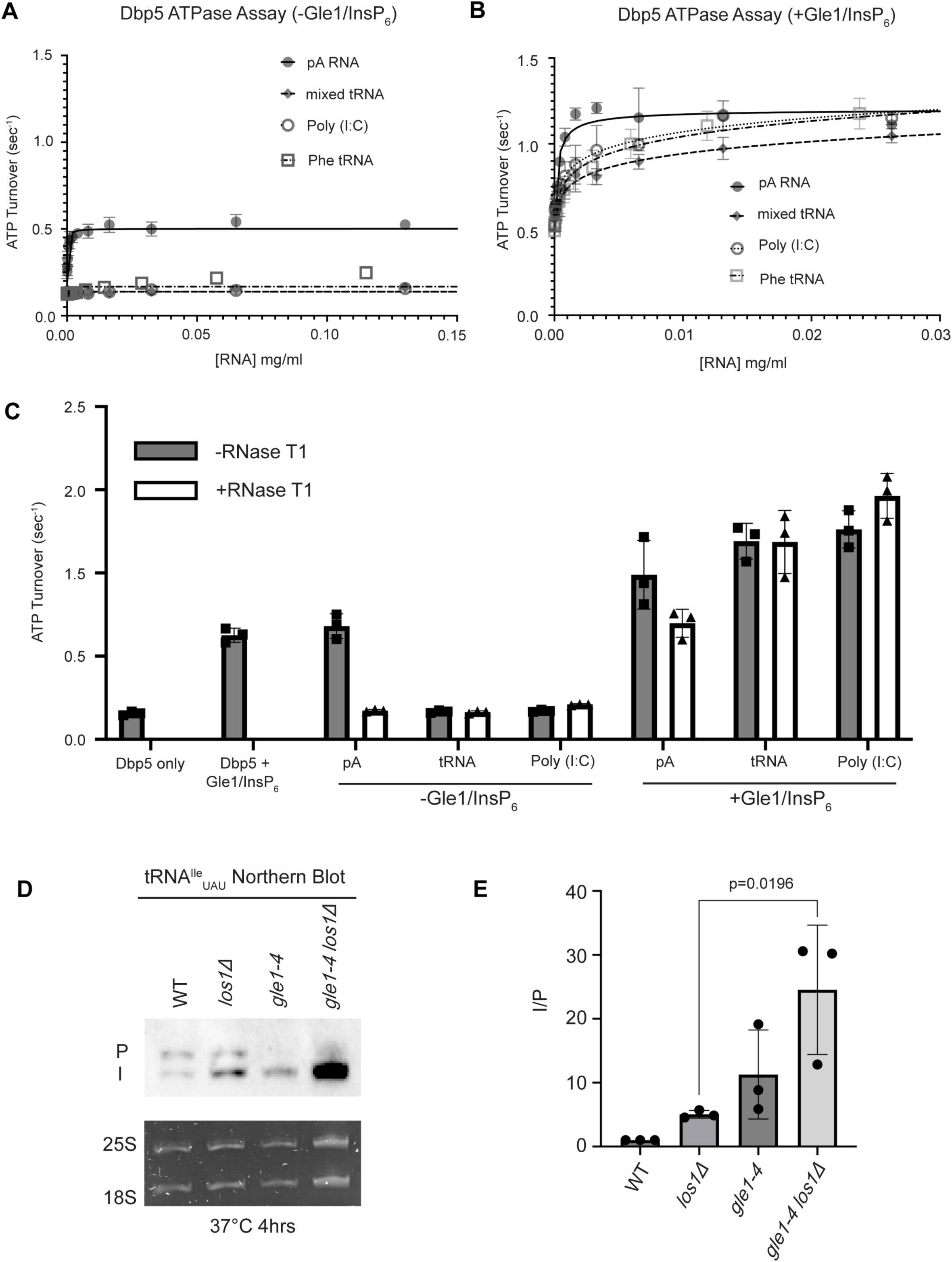
tRNA alone does not stimulate the Dbp5 ATPase cycle but can act synergistically with Gle1/InsP_6_ to fully activate Dbp5. (A) ATPase activity of Dbp5 is stimulated by ssRNA (pA, closed square) but not mixed yeast tRNA (closed diamond), Phe tRNA (open square), or poly (I:C) dsRNA (open circle). Steady-state ATPase assays were conducted in the presence of 1 µM Dbp5, 2.5 mM ATP, and varying concentrations of RNA. Data was fit to Michaelis-Menten equation using GraphPad Prism. Error bars represent standard deviation of three independent experiments. (B) Gle1/InsP_6_ synergistically stimulates Dbp5 ATPase activity with mixed yeast tRNA (closed diamond), Phe tRNA (open square) and poly (I:C) dsRNA (open circle) substrates like ssRNA (pA). Steady-state ATPase assays were conducted in presence of 1 µM Dbp5, 2.5 mM ATP, 2 µM Gle1, 2 µM InsP_6_, and varying concentration of RNA. Data was fit to an allosteric sigmoidal model in GraphPad Prism. Error bars represent standard deviation of three independent experiments. (C) RNase T1 treatment of RNA for 2 hours at 37°C prior to ATPase assays confirms that the observed synergistic activation of Dbp5 ATPase activity by Gle1/InsP_6_ and tRNA or dsRNA is not caused by low levels of contaminating ssRNA. Steady-state ATPase assays were conducted in presence of 1 µM Dbp5, 2.5 mM ATP, 0.2 mg/ml RNA and 2 µM Gle1/InsP_6_ when indicated. (D) Northern Blot analysis targeting precursor and mature isoforms of tRNA^Ile^_UAU_. Small RNAs were isolated from strains at mid log phase growth after pre-culture at 25°C and shift to 37°C for 4 hours. “P” bands represent intron containing precursors that have 5’ leader/3’ trailer sequences and “I” bands represent intron-containing end processed tRNA intermediates that have leader/trailer sequences removed. (E) Quantification of Northern blot from (D). Ratio of signal from intron-containing end processed intermediates (I) vs 5’ leader/3’ trailer containing precursor (P) was calculated and presented relative to I/P ratio observed for WT. Error bars represent standard deviation and p-values calculated using one-way ANOVA.

The nucleoporin Gle1 with the small molecule inositol hexakisphosphate (InsP_6_) has been shown to synergistically activate Dbp5 ATPase activity at low RNA concentrations (Weirich, Erzberger et al. 2006, Montpetit, Thomsen et al. 2011). These findings have led to a model of spatially regulated Dbp5 activity to promote mRNA-protein complex remodeling at the cytoplasmic face of an NPC where Gle1 is localized, resulting in directional mRNA export (Hodge, Colot et al. 1999, Strahm, Fahrenkrog et al. 1999, Alcazar-Roman, Tran et al. 2006, Weirich, Erzberger et al. 2006, Dossani, Weirich et al. 2009, Hodge, Tran et al. 2011, Montpetit, Thomsen et al. 2011, Noble, Tran et al. 2011, Adams, Mason et al. 2017, Wong, Gray et al. 2018, Arul Nambi Rajan and Montpetit 2021). In the context of tRNA, addition of Gle1/InsP_6_ maximally stimulated Dbp5 (1.03 +/- 0.04 ATP/s) to a level like that of ssRNA (1.11 +/- 0.07) (Figure 5B). The synergistic activation of Dbp5 ATPase activity by Gle1/InsP_6_ and tRNA is mirrored by poly(I:C) and observed with both mixed yeast tRNAs as well as yeast Phenylalanine tRNA (Figure 5B). To confirm that this enhanced RNA activation by Gle1/InsP_6_ was not the result of contaminating ssRNA, ATPase assays were performed after treatment of tRNA or poly (I:C) with RNase T1 for 2 hours at 37°C. RNase T1 degrades ssRNA, not dsRNA, allowing determination of whether RNA activation observed in the presence of Gle1/InsP_6_ is specific to properly folded tRNA and poly(I:C). As expected, treatment of the ssRNA (pA) with RNase T1 resulted in the loss of RNA-stimulated Dbp5 ATPase activity (0.68 +/- 0.06 before RNase T1 treatment to 0.17 +/- 0.01 after). Furthermore, RNase T1 treatment reduced pA/Gle1/InsP_6_ activation (0.98 +/- 0.17 before treatment to 0.69 +/-0.07 after) to that of Dbp5 and Gle1/InsP_6_ alone (0.62 +/- 0.03) (Figure 5C). However, RNase T1 treated tRNA remained unchanged in the ability to activate Dbp5 in the presence of Gle1/InsP_6_ (1.19+0.11 without vs 1.18+/-0.19 ATP/s with RNase T1 treatment). This was recapitulated with poly(I:C) (1.26 +/- 0.11 ATP/s without and 1.46 +/- 0.135 ATP/s with RNase T1), confirming that the synergistic activation of Dbp5 observed in the presence of Gle1/InsP_6_ is dependent on the structured tRNA or dsRNA. These findings provide for the possibility that Dbp5 engages tRNA in the cell without ATPase activation and could remain bound until the Dbp5-tRNA complex reaches Gle1 at the cytoplasmic side of a nuclear pore complex. In support of this model, a strong additive defect in pre- tRNA^Ile^_UAU_ processing was observed by Northern blotting when *los1Δ* was combined *gle1-4* (∼5-fold increase in the I/P ratio relative to *los1Δ* alone, Figure 5D/E) after a 4-hour incubation at 37°C. These data indicate that Gle1, like Dbp5, supports pre-tRNA export independent of Los1.

## DISCUSSION

The best characterized tRNA export factors in *S. cerevisiae*, Los1 and Msn5, are non-essential and the *los111/msn511* double mutant is viable despite the essential role of tRNA export (Takano, Endo et al. 2005, Murthi, Shaheen et al. 2010). These data suggest additional mechanisms for regulated tRNA export, which is supported by recent publications that have implicated mRNA and rRNA export factors such as Mex67, Nup159, Gle1, Dbp5, and Crm1 in tRNA export (Wu, Bao et al. 2015, Chatterjee, Majumder et al. 2017, Nostramo and Hopper 2020, Chatterjee, Marshall et al. 2022). However, a mechanistic understanding of how these factors function in tRNA export and how such roles differ or relate to previously characterized roles in RNA export is still unresolved. Here, a combination of *in vivo* and *in vitro* data provide evidence that (1) Dbp5 and Gle1 have functions independent of Los1 in tRNA export; (2) Dbp5 can bind tRNA directly *in vitro* and does so in a manner that does not require Los1, Msn5, and Mex67 *in vivo*; and (3) the Dbp5 ATPase cycle is uniquely modulated by Gle1/InsP_6_ in the presence of structured RNA (e.g. tRNA or dsRNA) *in vitro* and the ATPase cycle supports tRNA export *in vivo*.

DEAD-box proteins (including Dbp5) broadly interact with RNA via the phosphate backbone (Andersen, Ballut et al. 2006, Andersen, Ballut et al. 2006, Linder 2006, Sengoku, Nureki et al. 2006, Linder and Fuller-Pace 2013, Arul Nambi Rajan and Montpetit 2021). The observed Dbp5-tRNA interaction is nucleotide dependent, supporting a specificity to the observed binding (Figure 4A/B). Furthermore, structural data has elucidated the synergistic activation of Dbp5 ATPase cycle induced by Gle1/InsP_6_ and mRNA binding (Montpetit, Thomsen et al. 2011). While further structural analysis is critical for understanding the same synergistic ATPase activation observed with tRNA and Gle1/InsP_6_, these results also support direct and productive binding *in-vitro* with a structured RNA. The synergistic activation of Dbp5 by Poly (I:C) in the presence of Gle1/InsP_6_ suggests the interaction is sequence independent and is not driven by elements unique to tRNAs. Collectively, the *in vitro* characterization of Dbp5 and tRNA described here suggest two non-mutually exclusive possibilities for why Dbp5 is only activated by tRNA/dsRNA in the presence of Gle1/InsP_6_. First, it may be that Dbp5 binds tRNA to form an inhibited intermediate that is relieved by Gle1/InsP_6._ This is an attractive hypothesis that is unique from how Dbp5 engages ssRNA (e.g., mRNA) and potentially supports previously proposed export models that suggest Dbp5 may bind and travel with a tRNA from nucleus to cytoplasm (Lari, Arul Nambi Rajan et al. 2019). A second possibility is that Gle1/InsP_6_ enhances the affinity of Dbp5 for tRNA and “non-mRNA” substrates, which is supported by biochemical characterizations that show Gle1/InsP_6_ promotes a shift in the steady-state distribution of populated Dbp5 intermediates from weak to strong RNA binding states (Hodge, Tran et al. 2011, Montpetit, Thomsen et al. 2011, Noble, Tran et al. 2011, Wong, Cao et al. 2016, Wong, Gray et al. 2018, Gray, Cao et al. 2022). Future structural and biochemical studies are expected to distinguish among these possibilities. Moreover, Dbp5 was previously reported to not bind dsRNA substrates (Weirich, Erzberger et al. 2006), as such the findings that Dbp5 can bind, and in the presence of Gle1/InsP_6_, be activated by tRNA should motivate reevaluation of possible functions of Dbp5-Gle1/InsP_6_ in regulating other highly structured RNA or “non-canonical” substrates *in vivo*.

While it has been reported previously that tRNA fails to stimulate Dbp5 ATPase cycle (Tseng, Weaver et al. 1998), here it is shown that despite a lack of stimulation when Gle1/InsP_6_ is absent, Dbp5 is able to bind tRNA *in vitro* and *in vivo*. The ability of a tRNA to bind but not stimulate the ATPase cycle of Dbp5 has important implications for mechanisms by which Dbp5 may function in tRNA export. For example, in mRNA export it is thought that Dbp5 functions are localized at NPCs (Derrer, Mancini et al. 2019, Lari, Arul Nambi Rajan et al. 2019, Adams and Wente 2020), where the Dbp5 ATPase cycle is tightly regulated by co-factors Nup159 and Gle1/InsP_6_ (Hodge, Colot et al. 1999, Rollenhagen, Hodge et al. 2004, Hodge, Tran et al. 2011, Montpetit, Thomsen et al. 2011, Noble, Tran et al. 2011, Wong, Gray et al. 2018). In fact, recent studies have shown the essential function of both Dbp5 and Mex67 in mRNA export can be accomplished when these proteins are fused to nuclear pore components, in addition Dbp5 does not form a complex with mRNAs in the nucleus (Derrer, Mancini et al. 2019, Adams and Wente 2020). These reports suggest that Dbp5 does not travel with an mRNA from the nucleoplasm to the cytoplasmic face of an NPC as part of an mRNA-protein complex, rather Dbp5 likely engages mRNAs after they exit the NPC transport channel to direct the final step(s) of mRNA export (Figure 6). However, in contrast to this mRNA export model, the unique ability of Dbp5 to bind tRNA without ATP stimulation, and then be fully activated by Gle1/InsP_6_, presents the potential for a novel mechanism of function in regulating tRNA export. Namely, that Dbp5 may enter the nucleus to stably engage tRNA and direct events leading to export, following which Dbp5 is removed from the tRNA by Gle1/InsP_6_ upon nuclear exit (Figure 6). Importantly, in both mRNA and tRNA export, Gle1/InsP_6_ activation is required to stimulate ATPase activity and recycle the enzyme. In contrast, Dbp5 ATPase activity and Gle1 have been shown to be dispensable for pre-ribosomal subunit transport (Neumann, Wu et al. 2016). This raises interesting questions of whether RNA binding and nucleocytoplasmic shuttling of Dbp5 alone can achieve this function, where in the cell these interactions occur, and how rRNA substrates modulate Dbp5 activity.

**Figure 6:**
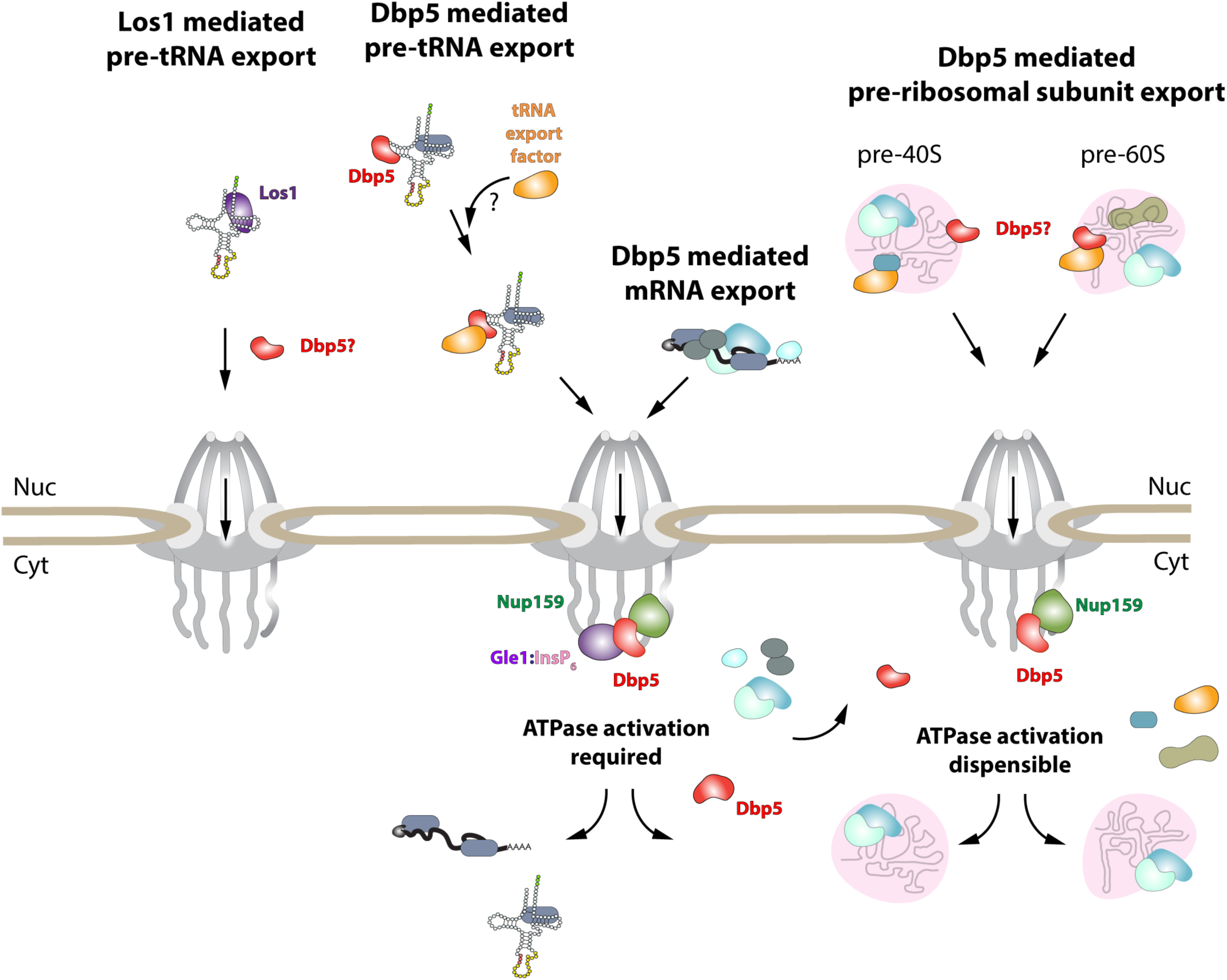
Model of Dbp5 function in mRNA, tRNA and pre-ribosomal subunit export. Data from this study support a model in which Dbp5-mediated export of both mRNA and tRNA require Gle1/InsP_6_ activation to remodel RNPs at the cytoplasmic face of the Nuclear Pore Complex. For tRNA export, lack of RNA-mediated ATPase stimulation following RNA binding may lead to formation of tRNA bound intermediates that act as adapters for recruitment of yet to be identified transport factors. For mRNA export, it has been shown that Dbp5 function is limited to the nuclear periphery and Dbp5 does not form stable nuclear complexes with mRNA. Gle1/InsP_6_ mediated activation of Dbp5 catalytic cycle and RNA release likely promotes recycling of export factors and Dbp5 to function in further rounds of export or other functions. In contrast, for Dbp5 mediated pre-ribosomal subunit export Dbp5 ATPase cycle and Gle1/InsP_6_ stimulation are dispensable for transport. As such, RNA binding may promote export through an unknown mechanism and that Dbp5-mediated remodeling does not occur at NPCs, with factors such as Mex67 persisting on pre-ribosomal subunits following export to the cytoplasm. While a role for Dbp5 functioning in the Los1 mediated pre-tRNA export pathway cannot be excluded, the data presented in this study support a Dbp5 mediated tRNA export pathway that exists independent of and parallel to Los1.

In mRNA export, it has been proposed that Dbp5 displaces mRNA export factors such as Mex67 and Nab2 to promote directionality of mRNA transit (Lund and Guthrie 2005, Tran, Zhou et al. 2007). Observations that Mex67 and Dbp5 appear to both function parallel to Los1 to promote tRNA export may point to an overlapping pathway that shares both mRNA export factors. Dbp5 and Mex67 have been proposed to form physical interactions, with Mex67 serving as an adaptor to direct Dbp5 to appropriate mRNA sites for remodeling (Lund and Guthrie 2005). However, here it is demonstrated that Mex67 is not required for Dbp5-tRNA interaction based on RNA IP assays. While Dbp5 recruitment to tRNA does not require Mex67, the data presented in this study do not exclude the possibility that the two proteins co-function to promote tRNA export. Indeed, previously published data suggests Mex67 likely requires an unknown adaptor to form its reported interactions with its tRNA substrates (Yao, Roser et al. 2007). Given the *in vitro* properties of Dbp5 binding to tRNA, it is tempting to speculate that nuclear Dbp5 promotes recruitment of Mex67 or Crm1 to tRNA-protein complexes. Indeed, a similar phenomenon has been described for the exon junction complex (EJC) and the DEAD-box protein at its core, eIF4AIII, that acts as a platform for the binding of other proteins to an mRNA (Ballut, Marchadier et al. 2005, Andersen, Ballut et al. 2006, Bono, Ebert et al. 2006, Pacheco-Fiallos, Vorlander et al. 2023). Stable binding of the EJC is governed by trans acting proteins that lock the complex on the mRNA by stabilizing an ADP-P_i_ hydrolysis immediate (Ballut, Marchadier et al. 2005, Bono, Ebert et al. 2006, Nielsen, Chamieh et al. 2009, Linder and Fuller-Pace 2013). In the case of Dbp5, the *in vitro* data suggests an inhibited state may be achieved in the context of tRNA alone, which could be further stabilized *in vivo* by yet to be discovered protein factors. The ability of Gle1/InsP_6_ to induce maximal activity of Dbp5 in the presence of tRNA would then allow for resolution of stable Dbp5-tRNA complexes in a spatially regulated manner to terminate export and recycle bound export factors (Figure 6). Alternatively, given that it has been reported that Mex67 can bind 5’ extended tRNAs and that these transcripts have been previously reported to contain 5’ methyl-guanosine cap structures (Ohira and Suzuki 2016, Chatterjee, Marshall et al. 2022), perhaps an “mRNA export-like” pathway that involves Cap Binding Complex (CBC) can act to recruit Mex67 to a subset of tRNAs. It remains unclear if CBC interacts with such transcripts and whether other export factors like Dbp5 also support “premature” export of unprocessed tRNAs.

More broadly, like many DEAD-box proteins, Dbp5 has many reported roles in controlling gene expression (Estruch and Cole 2003, Rollenhagen, Hodge et al. 2004, Gross, Siepmann et al. 2007, Scarcelli, Viggiano et al. 2008, Tieg and Krebber 2013, Wu, Becker et al. 2014, Neumann, Wu et al. 2016, Mikhailova, Shuvalova et al. 2017, Lari, Arul Nambi Rajan et al. 2019, Beissel, Grosse et al. 2020). The nucleic acid substrate-specific biochemical properties of Dbp5 ATPase regulation defined here may have wide-ranging implications for Dbp5 functions in pre-ribosomal RNA export, translation, or R-loop metabolism. Furthermore, several DEAD-box proteins such as Dbp2 have been reported to bind diverse nucleic acid substrates (ncRNA and mRNA), including G-Quadraplexes, and have roles in R-loop biology (Ma, Paudel et al. 2016, Xing, Wang et al. 2017, Tedeschi, Cloutier et al. 2018, Gao, Byrd et al. 2019, Lai, Choudhary et al. 2019, Lai and Tran 2021, Yan, Obi et al. 2021, Song, Lai et al. 2022). Given the high degree of conservation in the structure, function, and regulation of Dbp5 with other DEAD-box proteins (Ozgur, Buchwald et al. 2015, Sloan and Bohnsack 2018), the observations reported here raise the possibility that other DEAD-box proteins may also demonstrate substrate-specific regulation. Furthermore, the results here support the importance of Gle1 and the small molecule InsP_6_ as potent regulators of Dbp5 activity in multiple RNA export pathways. Thus, future investigation of how Dbp5 mediated export is modulated by regulation of these cellular factors during stress or disease contexts represent important areas of future study.

Overall, this study and other recent publications support a general function for both Dbp5 and Mex67 in RNA export (Yao, Roser et al. 2007, Yao, Lutzmann et al. 2008, Faza, Chang et al. 2012, Wu, Becker et al. 2014, Neumann, Wu et al. 2016, Chatterjee, Majumder et al. 2017, Becker, Hirsch et al. 2019, Lari, Arul Nambi Rajan et al. 2019, Nostramo and Hopper 2020, Vasianovich, Bajon et al. 2020, Chatterjee, Marshall et al. 2022). While these factors have been most well characterized in mRNA export, it is now important to reconsider Dbp5 and Mex67 as general RNA export factors, which raises questions about possible co-regulation of mRNA and non-coding RNA export pathways to fine tune gene expression. To this end, the *in vitro*, biochemical, and genetic properties of Dbp5 reported in this study will inform future structure-function and mechanistic characterization of a Dbp5 mediated, and Gle1/InsP_6_ regulated, tRNA export pathway(s). For example, it will be important for future studies to address the composition of pathway specific export-competent tRNA-protein complexes and how Dbp5 contributes to changes in the architecture of such a complex as they move from nucleus to cytoplasm and back again.

## MATERIALS AND METHODS

### Yeast strains generation and growth conditions

A list of all yeast strains used in this study is provided in Table S1. Deletion mutants and C-terminal tagging was conducted by PCR based transformations and confirmed by colony PCR. All yeast transformations were conducted using previously published standard high-efficiency LiAc, ssDNA, PEG protocol (Gietz and Woods 2002). Yeast were cultured in YPD or synthetic complete (SC) media when indicated to mid log growth. For overexpression of Dbp5 ATPase mutants, untagged Dbp5 or *dbp5^R426Q^, dbp5^R369G^,* or *dbp5^E240Q^* were integrated at URA3 locus under control of pGAL promoter in a wild-type strain with Dbp5 expressed from its endogenous locus and promoter. pGAL expression was induced by first derepressing the promoter with growth in YP media with 2% raffinose overnight followed by a shift to 2% galactose containing media for 6 hours. Overexpression was halted by addition of 2% glucose and further incubation for 1 hour.

### Spot growth assay

Yeast were pre-cultured to saturation overnight, diluted, and 1:10 serial dilutions were made with the most concentrated wells representing a concentration of 0.25 OD/ml. 3 µL of each dilution was spotted on YPD and grown at indicated temperatures for 2 days.

### Live Cell Imaging

Imaging experiments were carried out using the confocal configuration of an Andor Dragonfly microscope equipped with an EMCCD camera driven by Fusion software (Andor) using a 60x oil immersion objective (Olympus, numerical aperture (N.A.) 1.4). Images were acquired from cells grown to mid log phase in SC media at 25°C and shifted to 37°C for indicated time or treated with DMSO or 500uM Auxin and 10 uM InsP6 for relevant experiments. Cells were immobilized in 384-well glass bottomed plates (VWR) that were pre-treated with Concanavalin A.

### tRNA extraction and Northern Blotting

Isolation of small RNAs and Northern blot experiments were performed as previously described (Wu, Huang et al. 2013). Briefly, small RNAs were isolated from yeast cultures grown to early log phase by addition of equal volumes cold TSE buffer (0.01M Tris pH7.5, 0.01M EDTA, 0.1M sodium chloride) and TSE saturated phenol. Samples were incubated at 55°C for 20min with vortexing every 3 min then placed on ice for 10 min. Aqueous phase was extracted after centrifugation, re-extraction with phenol was performed, and RNA was precipitated overnight in ethanol at -80°C. 2.5ug of RNA was then separated on 10% TBE-Urea gels, gels were stained with Apex safe DNA gel stain to visualize 25S and 18S rRNA and then transferred to Hybond N^+^ membrane (Amersham). RNA was crosslinked to membranes at 2400J/m^2^ using UV Crosslinker (VWR) and tRNA were detected using digoxigenin-labeled (DIG) probes. Mean integrated intensity values were measured for I and P bands using FIJI (Schindelin, Arganda-Carreras et al. 2012) and normalized to background signal in each lane.

Sequence of Northern blot probe targeting precursor and mature isoforms of tRNA^Ile^_UAU_ used in Figure 2 and 4 is as follows: GGCACAGAAACTTCGGAAACCGAATGTTGCTATAAGCACGAAGCTCTAACCACTGAGCTACACGAGC.

### Fluorescence In-Situ Hybridization (FISH)

FISH experiments were carried out as described in previous publications with minor modifications (Lord, Ospovat et al. 2017) using directly Cy3 labeled tRNA probes or fluorescein isothiocyanate (FITC) labeled dT probes. Sequence of Cy3 labeled tRNA^Ile^_UAU_ intron specific probe is same as previously published SRIM03 (Chatterjee, Majumder et al. 2017) with Cy3 moiety appended at the 5’ end (CGTTGCTTTTAAAGGCCTGTTTGAAAGGTCTTTGGCACAGAAACTTCGGAAACCGAATGTTGCTAT). Briefly, yeast cultures were grown to early log phase at 25°C and treated with drug/vehicle or shifted to 37°C for 4 hours when appropriate. Samples were fixed with 0.1 volume of 37% formaldehyde for 15min. These samples were then pelleted and resuspended in 4% paraformaldehyde (PFA) solution (4% PFA, 0.1M Potassium Phosphate (pH 6.5), 0.5mM MgCl_2_) and incubated with rotation at 25°C for 3 hours. Cells were then washed twice with Buffer B (1.2M Sorbitol, 0.1M Potassium Phosphate (pH 6.5), 0.5mM MgCl_2_). For spheroplasting cell pellets were resuspended in a 1ml Buffer B solution containing 0.05% beta-mercaptoethanol. 15ul of 20mg/mL zymolyase T20 was then added to each sample and were subsequently incubated at 37°C for 45min. After washing once with Buffer B, cells were adhered to eight well slides (Fisher Scientific) pre-coated with Poly-L-Lysine. Slides were placed sequentially in 70%, 90%, and 100% ethanol for 5min and then allowed to dry. Samples were next incubated at 37°C for 2 hours in pre-hybridization buffer (4X SSC, 50% formamide, 10% dextran sulfate, 125 ug/mL *E. coli* tRNA, 500 ug/mL salmon sperm DNA, 1x Denhardt’s solution, and 10mM vanadyl ribonucleoside complex (VRC)). Pre-hybridization solution was then aspirated and replaced with pre-warmed pre-hybridization solution that contained either 0.25uM Cy3 tRNA probe or 0.025uM FITC dT probe and further incubated at 37°C overnight. Wells were then sequentially washed for 5min once with 2x SSC, three times with 1xSSC, and once with 0.5x SSC. Slides were dipped in 100% ethanol, air-dried, and mounting media with 4’, 6-diamidino-2-phenylidole (DAPI) was applied to each well prior to sealing of coverslip onto slide. Imaging was performed using Andor Dragonfly equipped with Andor iXon Ultra 888 EMCCD camera driven by Fusion software (Andor) using a 60x 1.4 N.A. oil objective. Images were processed in FIJI (e.g., cropping, brightness/contrast adjustments and maximum z-projections) (Schindelin, Arganda-Carreras et al. 2012). Quantification of tRNA FISH was performed by acquiring average nuclear and cytoplasmic pixel intensities respectively. Maximum projection images were generated from z stack images for DAPI and tRNA FISH channels. Nuclear and whole cell masks were then respectively generated using Cellpose software (Stringer, Wang et al. 2021, Pachitariu and Stringer 2022). Nuclear mask regions were removed from whole cell masks to generate cytoplasmic mask and average pixel intensities were calculated. The ratio of average nuclear and cytoplasmic mask pixel intensity per cell was calculated.

### RNA Immunoprecipitation (RIP)

RIPs were conducted as previously described with minor modifications (Lari, Arul Nambi Rajan et al. 2019). Pull downs were performed targeting protein-A (prA) tagged Dbp5 in parallel with an untagged control to assess background non-specific binding. Crosslinking was conducted on yeast cultures in mid-log growth by addition of formaldehyde to final concentration of 0.3% and incubation for 30 min. Crosslinking was quenched by addition of glycine to a final concentration of 60mM for 10 min. Cells were harvested and pellets frozen in liquid nitrogen. Lysis was performed using ice cold TN150 (50mM Tris-HCl pH7.8, 150mM NaCl, 0.1% IGEPAL, 5mM beta-mercaptoethanol) supplemented with 10mM EDTA, 1ng Luciferase Spike-In RNA (Promega), and 1X protease inhibitor cocktail. 1ml zirconia beads (0.5mm) were used for disrupting cells by vortexing 5 times for 30 seconds followed by 1min on ice between each pulse. Lysate was pre-cleared by centrifugation at 4000xg 5min followed by 20min at 20000xg at 4°C. 1% pre-IP lysate was preserved as input RNA sample and additional pre-IP sample was reserved for western blotting. Remaining lysates were then diluted to 10ml with TN150 and incubated with IgG-conjugated magnetic dynabeads at 4°C for 30min with constant rotation. Immunoprecipitate was washed once with 1ml TN150, once with 1ml TN1000 (50mM Tris-HCl pH7.8, 1M NaCl 0.1% IGEPAL, 5mM beta-mercaptoethanol), and again once with 1ml TN150 each for 5min at 4°C. Beads from RIPs were then resuspended in 500ul proteinase K elution mix (50mM Tris-HCl pH 7.8, 50mM NaCl, 1mM EDTA, 0.5% SDS, 100ug proteinase K) in parallel with input samples and incubated at 50°C for 2 hours followed by 65°C for 1 hour to allow crosslink reversal. RNA was isolated by extraction with acidic Phenol:Chloroform:Isoamylalcohol (pH 4.3-4.7) and ethanol precipitation. Half volume of RNA was reverse transcribed using Superscript III using random priming according to manufacturer instructions and other half of RNA was retained for minus RT control. RNA was analyzed by qPCR using Power SYBR (Applied Biosystems) on an Applied Biosystems instrument. Target RNA abundance in RIP was normalized to abundance of target in input sample and relative fold enrichment was calculated by comparing signal in specific prA-Dbp5 RIPs to signal from untagged control. Standard curves were generated to test PCR efficiencies of each primer set used in this study. Primer sequences used are as follows:

tRNA^Ile^_UAU_ Unspliced Forward: GCTCGTGTAGCTCAGTGGTTAG

tRNA^Ile^_UAU_ Unspliced Reverse: CTTTTAAAGGCCTGTTTGAAAG FBA1

Forward: CGAAAACGCTGACAAGGAAG

FBA1 Reverse: TCTCAAAGCGATGTCACCAG

### Western Blotting

∼5 OD pellets were lysed in laemmli buffer and loaded onto 10% SDS PAGE. Protein transfer was performed onto nitrocellulose membrane using cold wet tank transfer protocol. Western blotting was performed with 1:5000 dilution of mouse monoclonal anti-DBP5 and 1:10000 dilution of mouse monoclonal anti-GAPDH (Thermo Fisher) primary antibody overnight at 4°C and 1:10000 anti-mouse DyeLight 650 secondary to detect protein.

### Protein Purification, ATPase assays and Fluorescence Polarization

Protein purification, *in vitro* ATPase assays and fluorescence polarization using full-length Dbp5 performed as described previously (Weirich, Erzberger et al. 2006, Montpetit, Thomsen et al. 2011, Montpetit, Seeliger et al. 2012). Commercially available Poly (I:C), mixed yeast tRNA and yeast Phenylalanine tRNA were obtained from Sigma for use in indicated assays. For RNase T1 ATPase assays, indicated RNA samples were treated with 33 units of RNase T1 for 2 hours at 37°C prior to experiment. Fluorescence polarization experiments were performed using a fluorescein (fl) labelled 16nt ssRNA (5’-fl-GGGUAAAAAAAAAAAA-3’) in the presence of ADP•BeF_3_ with increasing concentrations of unlabeled polyadenylic acid or yeast mixed tRNA titrated into the reactions. Fluorescence polarization and ATPase assays were assembled in the same ATPase buffer (30mM HEPES [pH 7.5], 100mM NaCl, and 2mM MgCl_2_). All ATPase assays with RNA titrations were fit to Michaelis Menten or allosteric sigmoidal models (when Gle1/InsP_6_ was added) in GraphPad Prism.

### RNA Electrophoretic Mobility Shift Assay (EMSA) and denaturing urea PAGE

Recombinant full length Dbp5 was purified as above and described previously(Montpetit, Thomsen et al. 2011, Montpetit, Seeliger et al. 2012). RNA binding and EMSA was performed according to previously published assay conditions (Yao, Roser et al. 2007). RNA binding reactions were assembled using binding buffer (20mM HEPES [pH7.4], 100 mM KCl, 10 mM NaCl, 4 mM MgCl_2_, 0.2 mM EDTA, 20% glycerol, 1 mM DTT, 0.5% NP-40) in the presence of indicated nucleotide and loaded on to 6% native polyacrylamide el (0.5 x TBE). Gels were stained with Apex gel stain to visualize RNA. Mean integrated intensity of bands were quantified in FIJI (Schindelin, Arganda-Carreras et al. 2012) and normalized to background signal for each respective well. Bound fraction was calculated and fit to a one-site binding equation in GraphPad Prism.

## ACKNOWLEDGEMENTS

We would like to acknowledge and thank all current and past members of the Montpetit lab for their aid and helpful discussions over the course of this work. Particularly we thank Rachel Montpetit for assistance with cloning and strain construction. We would also like to thank members of the Aitchison and Wozniak laboratories for their constructive feedback during this study. AANR was funded in part by the predoctoral Training Program in Molecular and Cellular Biology at UC Davis that is supported by an NIH T32 training grant (GM007377). RA was supported by a postdoctoral fellowship from Uehara memorial foundation and an overseas research fellowship from Japan Society for the Promotion of Science. Research was further supported by the National Institute of General Medical Sciences of the National Institutes of Health under Award Number R35GM145328. The content is solely the responsibility of the authors and does not necessarily represent the views of funding agencies.

---

**Figure 1 Supplement:**
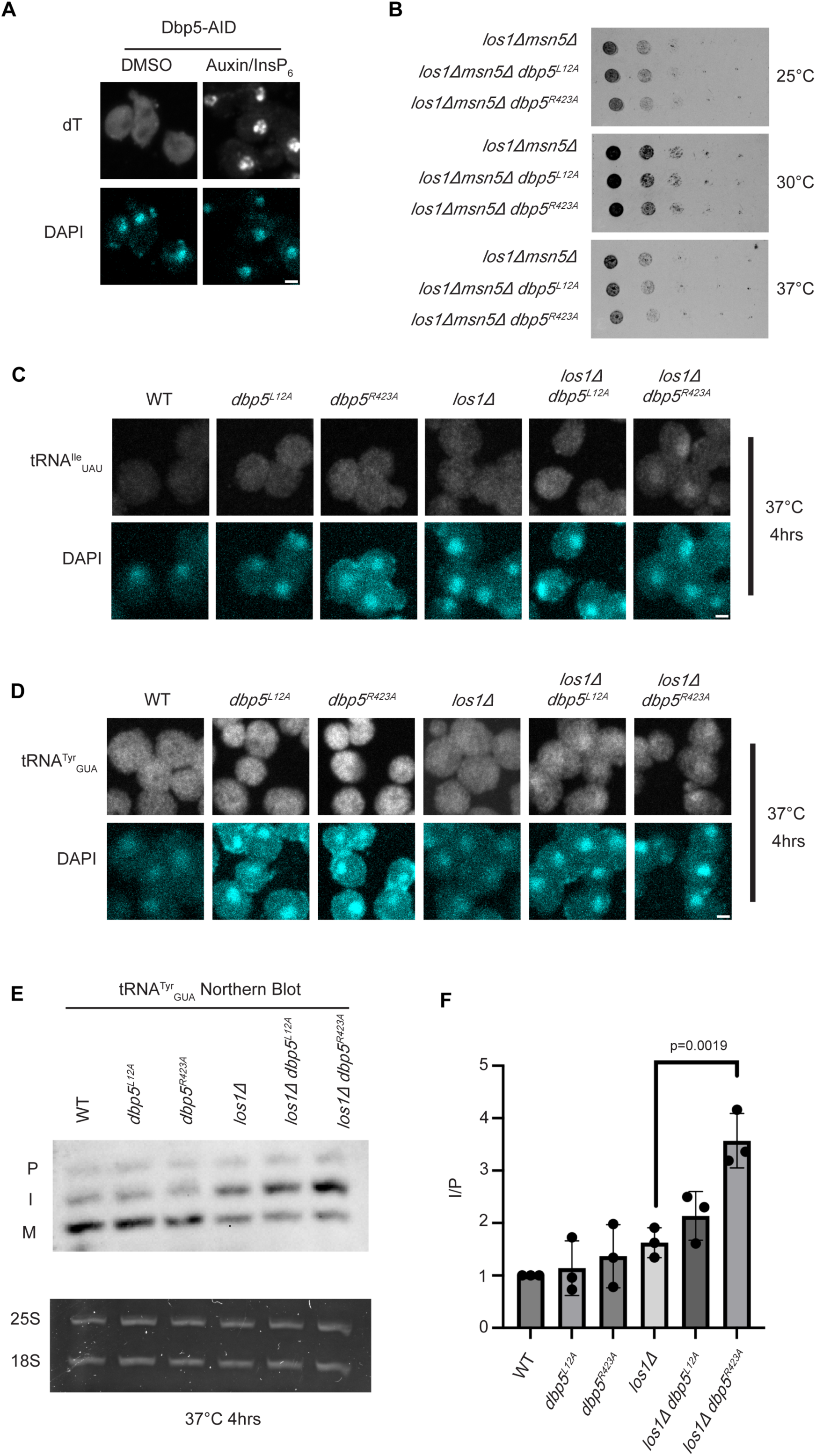
Dbp5 functions parallel to Los1 in pre-tRNA export. (A) dT FISH confirms induction of mRNA export defect and Dbp5 loss of function after addition of 500 µM Auxin and 10 µM InsP_6_ in Dbp5-AID strain for 90 minutes. Scale bar represents 2 µm. (B) Spot assay for growth of shuffle strains containing untagged *dbp5^L12A^* or *dbp5^R423A^* on Leu marked CEN plasmids in combination with *los111/msn511.* Genomic copy of Dbp5 has been replaced with His6Mx marker. Growth for two days at 25, 30, and 37°C on YPD was conducted following two rounds of counter selection for Ura marked WT Dbp5 CEN plasmids on 5’FOA. (C) tRNA FISH using a probe that can hybridize to the intron-containing and spliced isoform of tRNA^Ile^_UAU_ in indicated strains after pre-culture to early log phase at 25°C and shift to 37°C for 4 hours. Scale bar represents 2 µm. (D) tRNA FISH using a probe that can hybridize to the intron-containing and spliced isoform of tRNA^Tyr^_GUA_ in indicated strains after pre-culture to early log phase at 25°C and shift to 37°C for 4 hours. Scale bar represents 2 µm. (E) Northern Blot analysis targeting precursor and mature isoforms of tRNA^Tyr^_GUA_. Small RNAs were isolated from strains at mid log phase growth after pre-culture at 25°C and shift to 37°C for 4 hours. “P” bands represent intron containing precursors that have 5’ leader/3’ trailer sequences and “I” bands represent intron-containing end processed tRNA intermediates that have leader/trailer sequences removed. (F) Quantification of Northern blot from (E). Ratio of signal from intron-containing end processed intermediates (I) vs 5’ leader/3’ trailer containing precursor (P) was calculated and presented relative to I/P ratio observed for WT. Error bars represent standard deviation and p-values calculated using one-way ANOVA.

## Notes

### Competing Interest Statement

The authors have declared no competing interest.

### Summary of Updates

This version of the manuscript has been revised to include minor adjustments to the figures and texts, including improved images in Figure 1 and correction of figure labels.

## References

Adams, R. L., A. C. Mason, L. Glass, Aditi and S. R. Wente (2017). "Nup42 and IP(6) coordinate Gle1 stimulation of Dbp5/DDX19B for mRNA export in yeast and human cells." Traffic 18(12): 776–790.

Adams, R. L. and S. R. Wente (2020). "Dbp5 associates with RNA-bound Mex67 and Nab2 and its localization at the nuclear pore complex is sufficient for mRNP export and cell viability." PLoS Genet 16(10): e1009033.

Alcazar-Roman, A. R., E. J. Tran, S. Guo and S. R. Wente (2006). "Inositol hexakisphosphate and Gle1 activate the DEAD-box protein Dbp5 for nuclear mRNA export." Nat Cell Biol 8(7): 711–716.

Andersen, C. B., L. Ballut, J. S. Johansen, H. Chamieh, K. H. Nielsen, C. L. Oliveira, J. S. Pedersen, B. Seraphin, H. Le Hir and G. R. Andersen (2006). "Structure of the exon junction core complex with a trapped DEAD-box ATPase bound to RNA." Science 313(5795): 1968–1972.

Andersen, C. B., L. Ballut, J. S. Johansen, H. Chamieh, K. H. Nielsen, C. L. Oliveira, J. S. Pedersen, B. Séraphin, H. Le Hir and G. R. Andersen (2006). "Structure of the exon junction core complex with a trapped DEAD-box ATPase bound to RNA." Science 313(5795): 1968–1972.

Arul Nambi Rajan, A. and B. Montpetit (2021). "Emerging molecular functions and novel roles for the DEAD-box protein Dbp5/DDX19 in gene expression." Cell Mol Life Sci 78(5): 2019–2030.

Bai, B., H. M. Moore and M. Laiho (2013). "CRM1 and its ribosome export adaptor NMD3 localize to the nucleolus and affect rRNA synthesis." Nucleus 4(4): 315–325.

Ballut, L., B. Marchadier, A. Baguet, C. Tomasetto, B. Seraphin and H. Le Hir (2005). "The exon junction core complex is locked onto RNA by inhibition of eIF4AIII ATPase activity." Nat Struct Mol Biol 12(10): 861–869.

Becker, D., A. G. Hirsch, L. Bender, T. Lingner, G. Salinas and H. Krebber (2019). "Nuclear Pre-snRNA Export Is an Essential Quality Assurance Mechanism for Functional Spliceosomes." Cell Rep 27(11): 3199–3214 e3193.

Beissel, C., S. Grosse and H. Krebber (2020). "Dbp5/DDX19 between Translational Readthrough and Nonsense Mediated Decay." Int J Mol Sci 21(3).

Bono, F., J. Ebert, E. Lorentzen and E. Conti (2006). "The crystal structure of the exon junction complex reveals how it maintains a stable grip on mRNA." Cell 126(4): 713–725.

Chatterjee, K., S. Majumder, Y. Wan, V. Shah, J. Wu, H. Y. Huang and A. K. Hopper (2017). "Sharing the load: Mex67-Mtr2 cofunctions with Los1 in primary tRNA nuclear export." Genes Dev 31(21): 2186–2198.

Chatterjee, K., W. A. Marshall and A. K. Hopper (2022). "Three tRNA nuclear exporters in S. cerevisiae: parallel pathways, preferences, and precision." Nucleic Acids Res 50(17): 10140–10152.

Cook, A. G., N. Fukuhara, M. Jinek and E. Conti (2009). "Structures of the tRNA export factor in the nuclear and cytosolic states." Nature 461(7260): 60–65.

Derrer, C. P., R. Mancini, P. Vallotton, S. Huet, K. Weis and E. Dultz (2019). "The RNA export factor Mex67 functions as a mobile nucleoporin." J Cell Biol 218(12): 3967–3976.

Dossani, Z. Y., C. S. Weirich, J. P. Erzberger, J. M. Berger and K. Weis (2009). "Structure of the C-terminus of the mRNA export factor Dbp5 reveals the interaction surface for the ATPase activator Gle1." Proc Natl Acad Sci U S A 106(38): 16251–16256.

Estruch, F. and C. N. Cole (2003). "An early function during transcription for the yeast mRNA export factor Dbp5p/Rat8p suggested by its genetic and physical interactions with transcription factor IIH components." Mol Biol Cell 14(4): 1664–1676.

Eswara, M. B., A. T. McGuire, J. B. Pierce and D. Mangroo (2009). "Utp9p facilitates Msn5p-mediated nuclear reexport of retrograded tRNAs in Saccharomyces cerevisiae." Mol Biol Cell 20(23): 5007–5025.

Fan, J. S., Z. Cheng, J. Zhang, C. Noble, Z. Zhou, H. Song and D. Yang (2009). "Solution and crystal structures of mRNA exporter Dbp5p and its interaction with nucleotides." J Mol Biol 388(1): 1–10.

Faza, M. B., Y. Chang, L. Occhipinti, S. Kemmler and V. G. Panse (2012). "Role of Mex67-Mtr2 in the nuclear export of 40S pre-ribosomes." PLoS Genet 8(8): e1002915.

Feng, W. and A. K. Hopper (2002). "A Los1p-independent pathway for nuclear export of intronless tRNAs in Saccharomycescerevisiae." Proc Natl Acad Sci U S A 99(8): 5412–5417.

Gao, J., A. K. Byrd, B. L. Zybailov, J. C. Marecki, M. J. Guderyon, A. D. Edwards, S. Chib, K. L. West, Z. J. Waldrip, S. G. Mackintosh, Z. Gao, A. A. Putnam, E. Jankowsky and K. D. Raney (2019). "DEAD-box RNA helicases Dbp2, Ded1 and Mss116 bind to G-quadruplex nucleic acids and destabilize G-quadruplex RNA." Chem Commun (Camb) 55(31): 4467–4470.

Gietz, R. D. and R. A. Woods (2002). "Transformation of yeast by lithium acetate/single-stranded carrier DNA/polyethylene glycol method." Methods Enzymol 350: 87–96.

Graczyk, D., M. Ciesla and M. Boguta (2018). "Regulation of tRNA synthesis by the general transcription factors of RNA polymerase III - TFIIIB and TFIIIC, and by the MAF1 protein." Biochim Biophys Acta Gene Regul Mech 1861(4): 320–329.

Gray, S., W. Cao, B. Montpetit and E. M. De La Cruz (2022). "The nucleoporin Gle1 activates DEAD-box protein 5 (Dbp5) by promoting ATP binding and accelerating rate limiting phosphate release." Nucleic Acids Res 50(7): 3998–4011.

Gross, T., A. Siepmann, D. Sturm, M. Windgassen, J. J. Scarcelli, M. Seedorf, C. N. Cole and H. Krebber (2007). "The DEAD-box RNA helicase Dbp5 functions in translation termination." Science 315(5812): 646–649.

Guerrier-Takada, C., K. Gardiner, T. Marsh, N. Pace and S. Altman (1983). "The RNA moiety of ribonuclease P is the catalytic subunit of the enzyme." Cell 35(3 Pt 2): 849–857.

Harris, M. E. (2016). "Theme and Variation in tRNA 5’ End Processing Enzymes: Comparative Analysis of Protein versus Ribonucleoprotein RNase P." J Mol Biol 428(1): 5–9.

Haruki, H., J. Nishikawa and U. K. Laemmli (2008). "The anchor-away technique: rapid, conditional establishment of yeast mutant phenotypes." Mol Cell 31(6): 925–932.

Hodge, C. A., H. V. Colot, P. Stafford and C. N. Cole (1999). "Rat8p/Dbp5p is a shuttling transport factor that interacts with Rat7p/Nup159p and Gle1p and suppresses the mRNA export defect of xpo1-1 cells." EMBO J 18(20): 5778–5788.

Hodge, C. A., E. J. Tran, K. N. Noble, A. R. Alcazar-Roman, R. Ben-Yishay, J. J. Scarcelli, A. W. Folkmann, Y. Shav-Tal, S. R. Wente and C. N. Cole (2011). "The Dbp5 cycle at the nuclear pore complex during mRNA export I: dbp5 mutants with defects in RNA binding and ATP hydrolysis define key steps for Nup159 and Gle1." Genes & development 25(10): 1052–1064.

Hodge, C. A., E. J. Tran, K. N. Noble, A. R. Alcazar-Roman, R. Ben-Yishay, J. J. Scarcelli, A. W. Folkmann, Y. Shav-Tal, S. R. Wente and C. N. Cole (2011). "The Dbp5 cycle at the nuclear pore complex during mRNA export I: dbp5 mutants with defects in RNA binding and ATP hydrolysis define key steps for Nup159 and Gle1." Genes Dev 25(10): 1052–1064.

Hopper, A. K. and H. Y. Huang (2015). "Quality Control Pathways for Nucleus-Encoded Eukaryotic tRNA Biosynthesis and Subcellular Trafficking." Mol Cell Biol 35(12): 2052–2058.

Huang, H. Y. and A. K. Hopper (2015). "In vivo biochemical analyses reveal distinct roles of beta-importins and eEF1A in tRNA subcellular traffic." Genes Dev 29(7): 772–783.

Jankowsky, E. (2011). "RNA helicases at work: binding and rearranging." Trends Biochem Sci 36(1): 19–29.

Kaminski, T., J. P. Siebrasse and U. Kubitscheck (2013). "A single molecule view on Dbp5 and mRNA at the nuclear pore." Nucleus 4(1): 8–13.

Lai, Y. H., K. Choudhary, S. C. Cloutier, Z. Xing, S. Aviran and E. J. Tran (2019). "Genome-Wide Discovery of DEAD-Box RNA Helicase Targets Reveals RNA Structural Remodeling in Transcription Termination." Genetics 212(1): 153–174.

Lai, Y. H. and E. J. Tran (2021). "Probing Transcriptome-Wide RNA Structural Changes Dependent on the DEAD-box Helicase Dbp2." Methods Mol Biol 2209: 287–305.

Lari, A., A. Arul Nambi Rajan, R. Sandhu, T. Reiter, R. Montpetit, B. P. Young, C. J. Loewen and B. Montpetit (2019). "A nuclear role for the DEAD-box protein Dbp5 in tRNA export." Elife 8.

Linder, P. (2006). "Dead-box proteins: a family affair—active and passive players in RNP-remodeling." Nucleic acids research 34(15): 4168–4180.

Linder, P. and F. V. Fuller-Pace (2013). "Looking back on the birth of DEAD-box RNA helicases." Biochim Biophys Acta 1829(8): 750–755.

Linder, P. and F. V. Fuller-Pace (2013). "Looking back on the birth of DEAD-box RNA helicases." Biochimica et Biophysica Acta (BBA)-Gene Regulatory Mechanisms 1829(8): 750–755.

Lord, C. L., O. Ospovat and S. R. Wente (2017). "Nup100 regulates Saccharomyces cerevisiae replicative life span by mediating the nuclear export of specific tRNAs." RNA 23(3): 365–377.

Lund, M. K. and C. Guthrie (2005). "The DEAD-box protein Dbp5p is required to dissociate Mex67p from exported mRNPs at the nuclear rim." Mol Cell 20(4): 645–651.

Ma, W. K., B. P. Paudel, Z. Xing, I. G. Sabath, D. Rueda and E. J. Tran (2016). "Recruitment, Duplex Unwinding and Protein-Mediated Inhibition of the Dead-Box RNA Helicase Dbp2 at Actively Transcribed Chromatin." J Mol Biol 428(6): 1091–1106.

Mason, A. C. and S. R. Wente (2020). "Functions of Gle1 are governed by two distinct modes of self-association." J Biol Chem 295(49): 16813–16825.

Mikhailova, T., E. Shuvalova, A. Ivanov, D. Susorov, A. Shuvalov, P. M. Kolosov and E. Alkalaeva (2017). "RNA helicase DDX19 stabilizes ribosomal elongation and termination complexes." Nucleic Acids Res 45(3): 1307–1318.

Montpetit, B., M. A. Seeliger and K. Weis (2012). "Analysis of DEAD-box proteins in mRNA export." Methods Enzymol 511: 239–254.

Montpetit, B., N. D. Thomsen, K. J. Helmke, M. A. Seeliger, J. M. Berger and K. Weis (2011). "A conserved mechanism of DEAD-box ATPase activation by nucleoporins and InsP6 in mRNA export." Nature 472(7342): 238–242.

Montpetit, B., N. D. Thomsen, K. J. Helmke, M. A. Seeliger, J. M. Berger and K. Weis (2011). "A conserved mechanism of DEAD-box ATPase activation by nucleoporins and InsP 6 in mRNA export." Nature 472(7342): 238–242.

Murthi, A., H. H. Shaheen, H. Y. Huang, M. A. Preston, T. P. Lai, E. M. Phizicky and A. K. Hopper (2010). "Regulation of tRNA bidirectional nuclear-cytoplasmic trafficking in Saccharomyces cerevisiae." Mol Biol Cell 21(4): 639–649.

Neumann, B., H. Wu, A. Hackmann and H. Krebber (2016). "Nuclear Export of Pre-Ribosomal Subunits Requires Dbp5, but Not as an RNA-Helicase as for mRNA Export." PLoS One 11(2): e0149571.

Nielsen, K. H., H. Chamieh, C. B. Andersen, F. Fredslund, K. Hamborg, H. Le Hir and G. R. Andersen (2009). "Mechanism of ATP turnover inhibition in the EJC." RNA 15(1): 67–75.

Nishimura, K., T. Fukagawa, H. Takisawa, T. Kakimoto and M. Kanemaki (2009). "An auxin-based degron system for the rapid depletion of proteins in nonplant cells." Nat Methods 6(12): 917–922.

Noble, K. N., E. J. Tran, A. R. Alcazar-Roman, C. A. Hodge, C. N. Cole and S. R. Wente (2011). "The Dbp5 cycle at the nuclear pore complex during mRNA export II: nucleotide cycling and mRNP remodeling by Dbp5 are controlled by Nup159 and Gle1." Genes Dev 25(10): 1065–1077.

Nostramo, R. T. and A. K. Hopper (2020). "A novel assay provides insight into tRNAPhe retrograde nuclear import and re-export in S. cerevisiae." Nucleic Acids Res 48(20): 11577–11588.

Ohira, T. and T. Suzuki (2016). "Precursors of tRNAs are stabilized by methylguanosine cap structures." Nat Chem Biol 12(8): 648–655.

Ozgur, S., G. Buchwald, S. Falk, S. Chakrabarti, J. R. Prabu and E. Conti (2015). "The conformational plasticity of eukaryotic RNA-dependent ATPases." FEBS J 282(5): 850–863.

Pacheco-Fiallos, B., M. K. Vorlander, D. Riabov-Bassat, L. Fin, F. J. O’Reilly, F. I. Ayala, U. Schellhaas, J. Rappsilber and C. Plaschka (2023). "mRNA recognition and packaging by the human transcription-export complex." Nature 616(7958): 828–835.

Pachitariu, M. and C. Stringer (2022). "Cellpose 2.0: how to train your own model." Nat Methods 19(12): 1634–1641.

Phizicky, E. M. and A. K. Hopper (2023). "The Life and Times of a tRNA." RNA.

Rollenhagen, C., C. A. Hodge and C. N. Cole (2004). "The nuclear pore complex and the DEAD box protein Rat8p/Dbp5p have nonessential features which appear to facilitate mRNA export following heat shock." Mol Cell Biol 24(11): 4869–4879.

Scarcelli, J. J., S. Viggiano, C. A. Hodge, C. V. Heath, D. C. Amberg and C. N. Cole (2008). "Synthetic genetic array analysis in Saccharomyces cerevisiae provides evidence for an interaction between RAT8/DBP5 and genes encoding P-body components." Genetics 179(4): 1945–1955.

Schindelin, J., I. Arganda-Carreras, E. Frise, V. Kaynig, M. Longair, T. Pietzsch, S. Preibisch, C. Rueden, S. Saalfeld, B. Schmid, J. Y. Tinevez, D. J. White, V. Hartenstein, K. Eliceiri, P. Tomancak and A. Cardona (2012). "Fiji: an open-source platform for biological-image analysis." Nat Methods 9(7): 676–682.

Schmitt, C., C. von Kobbe, A. Bachi, N. Pante, J. P. Rodrigues, C. Boscheron, G. Rigaut, M. Wilm, B. Seraphin, M. Carmo-Fonseca and E. Izaurralde (1999). "Dbp5, a DEAD-box protein required for mRNA export, is recruited to the cytoplasmic fibrils of nuclear pore complex via a conserved interaction with CAN/Nup159p." EMBO J 18(15): 4332–4347.

Sengoku, T., O. Nureki, A. Nakamura, S. Kobayashi and S. Yokoyama (2006). "Structural basis for RNA unwinding by the DEAD-box protein Drosophila Vasa." Cell 125(2): 287–300.

Shi, H. and P. B. Moore (2000). "The crystal structure of yeast phenylalanine tRNA at 1.93 A resolution: a classic structure revisited." RNA 6(8): 1091–1105.

Skowronek, E., P. Grzechnik, B. Spath, A. Marchfelder and J. Kufel (2014). "tRNA 3’ processing in yeast involves tRNase Z, Rex1, and Rrp6." RNA 20(1): 115–130.

Sloan, K. E. and M. T. Bohnsack (2018). "Unravelling the Mechanisms of RNA Helicase Regulation." Trends Biochem Sci 43(4): 237–250.

Snay-Hodge, C. A., H. V. Colot, A. L. Goldstein and C. N. Cole (1998). "Dbp5p/Rat8p is a yeast nuclear pore-associated DEAD-box protein essential for RNA export." EMBO J 17(9): 2663–2676.

Song, Q. X., C. W. Lai, N. N. Liu, X. M. Hou and X. G. Xi (2022). "DEAD-box RNA helicase Dbp2 binds to G-quadruplex nucleic acids and regulates different conformation of G-quadruplex DNA." Biochem Biophys Res Commun 634: 182–188.

Steiner-Mosonyi, M., D. M. Leslie, H. Dehghani, J. D. Aitchison and D. Mangroo (2003). "Utp8p is an essential intranuclear component of the nuclear tRNA export machinery of Saccharomyces cerevisiae." J Biol Chem 278(34): 32236–32245.

Strahm, Y., B. Fahrenkrog, D. Zenklusen, E. Rychner, J. Kantor, M. Rosbach and F. Stutz (1999). "The RNA export factor Gle1p is located on the cytoplasmic fibrils of the NPC and physically interacts with the FG-nucleoporin Rip1p, the DEAD-box protein Rat8p/Dbp5p and a new protein Ymr 255p." EMBO J 18(20): 5761–5777.

Stringer, C., T. Wang, M. Michaelos and M. Pachitariu (2021). "Cellpose: a generalist algorithm for cellular segmentation." Nat Methods 18(1): 100–106.

Takano, A., T. Endo and T. Yoshihisa (2005). "tRNA actively shuttles between the nucleus and cytosol in yeast." Science 309(5731): 140–142.

Takemura, R., Y. Inoue and S. Izawa (2004). "Stress response in yeast mRNA export factor: reversible changes in Rat8p localization are caused by ethanol stress but not heat shock." J Cell Sci 117(Pt 18): 4189–4197.

Tedeschi, F. A., S. C. Cloutier, E. J. Tran and E. Jankowsky (2018). "The DEAD-box protein Dbp2p is linked to noncoding RNAs, the helicase Sen1p, and R-loops." RNA 24(12): 1693–1705.

Thoms, M., E. Thomson, J. Bassler, M. Gnadig, S. Griesel and E. Hurt (2015). "The Exosome Is Recruited to RNA Substrates through Specific Adaptor Proteins." Cell 162(5): 1029–1038.

Tieg, B. and H. Krebber (2013). "Dbp5 - from nuclear export to translation." Biochim Biophys Acta 1829(8): 791–798.

Tran, E. J., Y. Zhou, A. H. Corbett and S. R. Wente (2007). "The DEAD-box protein Dbp5 controls mRNA export by triggering specific RNA:protein remodeling events." Mol Cell 28(5): 850–859.

Trotta, C. R., F. Miao, E. A. Arn, S. W. Stevens, C. K. Ho, R. Rauhut and J. N. Abelson (1997). "The yeast tRNA splicing endonuclease: a tetrameric enzyme with two active site subunits homologous to the archaeal tRNA endonucleases." Cell 89(6): 849–858.

Tseng, S. S., P. L. Weaver, Y. Liu, M. Hitomi, A. M. Tartakoff and T. H. Chang (1998). "Dbp5p, a cytosolic RNA helicase, is required for poly(A)+ RNA export." EMBO J 17(9): 2651–2662.

Tudek, A., M. Schmid, M. Makaras, J. D. Barrass, J. D. Beggs and T. H. Jensen (2018). "A Nuclear Export Block Triggers the Decay of Newly Synthesized Polyadenylated RNA." Cell Rep 24(9): 2457–2467 e2457.

Vasianovich, Y., E. Bajon and R. J. Wellinger (2020). "Telomerase biogenesis requires a novel Mex67 function and a cytoplasmic association with the Sm(7) complex." Elife 9.

von Moeller, H., C. Basquin and E. Conti (2009). "The mRNA export protein DBP5 binds RNA and the cytoplasmic nucleoporin NUP214 in a mutually exclusive manner." Nat Struct Mol Biol 16(3): 247–254.

Wan, Y. and A. K. Hopper (2018). "From powerhouse to processing plant: conserved roles of mitochondrial outer membrane proteins in tRNA splicing." Genes Dev 32(19-20): 1309–1314.

Weirich, C. S., J. P. Erzberger, J. M. Berger and K. Weis (2004). "The N-terminal domain of Nup159 forms a beta-propeller that functions in mRNA export by tethering the helicase Dbp5 to the nuclear pore." Mol Cell 16(5): 749–760.

Weirich, C. S., J. P. Erzberger, J. S. Flick, J. M. Berger, J. Thorner and K. Weis (2006). "Activation of the DExD/H-box protein Dbp5 by the nuclear-pore protein Gle1 and its coactivator InsP6 is required for mRNA export." Nat Cell Biol 8(7): 668–676.

Wong, E. V., W. Cao, J. Voros, M. Merchant, Y. Modis, D. D. Hackney, B. Montpetit and E. M. De La Cruz (2016). "P(I) Release Limits the Intrinsic and RNA-Stimulated ATPase Cycles of DEAD-Box Protein 5 (Dbp5)." J Mol Biol 428(2 Pt B): 492–508.

Wong, E. V., S. Gray, W. Cao, R. Montpetit, B. Montpetit and E. M. De La Cruz (2018). "Nup159 Weakens Gle1 Binding to Dbp5 But Does Not Accelerate ADP Release." J Mol Biol 430(14): 2080–2095.

Wu, H., D. Becker and H. Krebber (2014). "Telomerase RNA TLC1 shuttling to the cytoplasm requires mRNA export factors and is important for telomere maintenance." Cell Rep 8(6): 1630–1638.

Wu, J., A. Bao, K. Chatterjee, Y. Wan and A. K. Hopper (2015). "Genome-wide screen uncovers novel pathways for tRNA processing and nuclear-cytoplasmic dynamics." Genes Dev 29(24): 2633–2644.

Wu, J. and A. K. Hopper (2014). "Healing for destruction: tRNA intron degradation in yeast is a two-step cytoplasmic process catalyzed by tRNA ligase Rlg1 and 5’-to-3’ exonuclease Xrn1." Genes Dev 28(14): 1556–1561.

Wu, J., H. Y. Huang and A. K. Hopper (2013). "A rapid and sensitive non-radioactive method applicable for genome-wide analysis of Saccharomyces cerevisiae genes involved in small RNA biology." Yeast 30(4): 119–128.

Xing, Z., S. Wang and E. J. Tran (2017). "Characterization of the mammalian DEAD-box protein DDX5 reveals functional conservation with S. cerevisiae ortholog Dbp2 in transcriptional control and glucose metabolism." RNA 23(7): 1125–1138.

Yan, K. K., I. Obi and N. Sabouri (2021). "The RGG domain in the C-terminus of the DEAD box helicases Dbp2 and Ded1 is necessary for G-quadruplex destabilization." Nucleic Acids Res 49(14): 8339–8354.

Yao, W., M. Lutzmann and E. Hurt (2008). "A versatile interaction platform on the Mex67-Mtr2 receptor creates an overlap between mRNA and ribosome export." EMBO J 27(1): 6–16.

Yao, W., D. Roser, A. Kohler, B. Bradatsch, J. Bassler and E. Hurt (2007). "Nuclear export of ribosomal 60S subunits by the general mRNA export receptor Mex67-Mtr2." Mol Cell 26(1): 51–62.

Yoshihisa, T., C. Ohshima, K. Yunoki-Esaki and T. Endo (2007). "Cytoplasmic splicing of tRNA in Saccharomyces cerevisiae." Genes Cells 12(3): 285–297.

Yoshihisa, T., K. Yunoki-Esaki, C. Ohshima, N. Tanaka and T. Endo (2003). "Possibility of cytoplasmic pre-tRNA splicing: the yeast tRNA splicing endonuclease mainly localizes on the mitochondria." Mol Biol Cell 14(8): 3266–3279.

Zhao, J., S. B. Jin, B. Bjorkroth, L. Wieslander and B. Daneholt (2002). "The mRNA export factor Dbp5 is associated with Balbiani ring mRNP from gene to cytoplasm." EMBO J 21(5): 1177–1187.

